# Intrinsic Lipid Curvatures of Mammalian Plasma Membrane Outer Leaflet Lipids and Ceramides

**DOI:** 10.1101/2021.04.26.441390

**Authors:** Michael Kaltenegger, Johannes Kremser, Moritz P. Frewein, Primož Ziherl, Douwe J. Bonthuis, Georg Pabst

**Author notes:** these authors have contributed equally to the present work (Michael Kaltenegger, Johannes Kremser).

## Abstract

We developed a global X-ray data analysis method to determine the intrinsic curvatures of lipids hosted in inverted hexagonal phases. In particular, we combined compositional modelling with molecular shape-based arguments to account for non-linear mixing effects of guest-in-host lipids on intrinsic curvature. The technique was verified by all-atom molecular dynamics simulations and applied to sphingomyelin and a series of phosphatidylcholines and ceramides with differing composition of the hydrocarbon chains. We report positive lipid curvatures for sphingomyelin and all phosphatidylcholines with disaturated and monounsaturated hydrocarbons. Substitution of the second saturated hydrocarbon with an unsaturated acyl chain in turn shifted the intrinsic lipid curvatures to negative values. All ceramides, with chain lengths varying between C2:0 and C24:0, displayed significant negative lipid curvature values. Moreover, we report non-additive mixing for C2:0 ceramide and sphingomyelin. Our findings manifest the high and manifold potential of lipids to modulate physiological membrane function.

**Highlights:**

- Molecular shape based theory for non-linear lipid curvature mixing
- Global analysis of SAXS patterns inverted hexagonal phases using compositional modelling and Bayesian probability theory
- MD simulations of inverted hexagonal phases
- Non-additive mixing of palmitoyl sphingomyelin and C2:0 ceramide

## 1. Introduction

Membrane curvature is well-recognized for its key role in diverse cellular processes related to membrane remodelling, including fusion and fission [1, 2, 3], as well as tissue formation [4]. Intriguingly, curvature also plays a fundamental role in flat membranes. The stored elastic curvature stress, resulting from oppositely curved lipid leaflets brought together to form a flat bilayer may couple for example to the conformational equilibrium of integral or peripheral proteins [5, 6, 7]. Additionally, membrane curvature stress has been related to the activity of membrane-active compounds [8], stability of lipid asymmetry in plasma membranes [9], and membrane raft formation [10].

Leaflet curvature, often referred to as monolayer spontaneous curvature or intrinsic (spontaneous) lipid curvature and denoted by *C*_0_, is a result of the effective shape of its constituent molecules. That is, its value depends on the lipids’ steric size, as well as on the attractive and repulsive interactions along the molecular axis. Shape-based packing arguments in general have provided simple, but overall good understanding of thermodynamic stability, preferred size, and size distribution of lipid aggregates [11].

We here explore such shape arguments, which must, due to their nonspecific nature, apply equally well to lipid mixtures and to single-component aggregates. In particular, we apply packing arguments to derive the intrinsic curvatures of lipids that do not form inverted hexagonal phases (H_II_) in aqueous solution. Inverted hexagonal mesophases are convenient for the determination of *C*_0_ using structure sensitive techniques, such as small-angle X-ray scattering (SAXS) [12]. In order to derive *C*_0_ for bilayer-forming lipids, these lipids (‘guest’) are commonly added to H_II_-forming templates (‘host’) assuming linear mixing [13, 14, 15, 16, 17, 18]. Yet, May and Ben-Shaul pointed out almost three decades ago – using either mean-field molecular theory, or an elastic deformation model – that *C*_0_ may exhibit non-linear mixing contributions, given significant differences in chain length between host and guest lipids [19]. However, the involved model parameters evade experimental determination. Applying shape arguments for lipid mixing instead allowed us to derive a simple model for nonlinear mixing of lipids in H_II_ phases that can be exploited experimentally. We also alude to the possibility of non-additive mixing, leading to a variation of *C*_0_ of a lipids with its concentration, which has been implied to occur for cholesterol and sphingomyelin [20].

To determine *C*_0_, we applied SAXS using dioleoyl phosphatidylethanolamine (DOPE) as host lipid. DOPE forms a H_II_ phase over a broad range of temperatures [21] and has been used by several groups for such measurements [13, 14, 15, 16, 17, 18]. As guest lipids we studied phosphatidylcholines (PCs), palmitoyl sphingomyelin (PSM), as well as a series of ceramides (Cer) with different hydrocarbon chain lengths. PCs and sphinomyelin are both abundant in the outer leaflet of mammalian plasma membranes [22]. Ceramides in turn, except for the skin stratum corneum, occur only in trace amounts in cellular membranes (see [23], for a recent comprehensive review). Ceramides, due to their high hydrophobicity and multiple hydrogen bond formation abilities, significantly affect the physical properties of membranes and induce either the formation of highly-ordered domains, or lead to lipid scrambling (flip-flop) [23]. Of specific interest for the present study is their ability to promote H_II_ phase formation, indicating significant negative *C*_0_-values [24, 25].

We previously reported intrinsic lipid curvature values for a series of outer leaflet lipids [17]. However, the applied technique led, in particular for cylinder-shaped lipids such as PCs with saturated hydrocarbons and sphingomyelin, to significant experimental uncertainties due to their low solubility in H_II_ phases. We also reported a *C*_0_ similar to that of DOPE for C16:0 ceramide [26]. However, basing the analysis on shifts in Bragg-peak positions only might have led to large deviations from the true value of the intrinsic curvature.

In order to address these issues, we advance a previously reported compositional analysis technique for H_II_ phases [12] to lipid mixtures including non-linear mixing contributions based on lipid shape. The model is validated against all-atom molecular dynamics simulations and then applied to PCs with saturated, unsaturated and branched hydrocarbons, PSM, as well as C2:0, C6:0, C16:0 and C24:0 ceramide. In general we find a high sensitivity of our method to lipid headgroup and hydrocarbon chain structure. We derive slightly positive curvatures for all PCs with saturated chains increasing moderately toward shorted chain lengths. Inducing double bonds or additional methyl groups in the hydrocarbon regime in turn progressively shifted the intrinsic lipid curvatures to negative values. From the studied sphingolipids only PSM exhibits a clear positive curvature and indicated potential non-additive mixing, albeit no conclusive answer can be given due to the limited achievable PSM concentration range in DOPE mixtures. Finally, while all ceramides exhibited significant negative curvatures in general, we find pronounced non-additive mixing for C2:0 ceramide.

The paper is structured as follows: After the materials and methods section we introduce the concept of non-linear mixing based on shape arguments and detail how this is assessed by compositional SAXS data modelling including an optimization procedure based on Bayesian probability theory. We then test the modelling against all atom MD simulations of H_II_ phases and previously reported *C*_0_’s of DOPE and dipalmitoleoyl phosphatidylethanolamine (16:1 PE). Finally, the remainder of the results section provides details of non-linear and non-additive mixing contributions to intrinsic curvatures of the above mentioned glycerophospholipids and sphingolipids, which are discussed in the subsequent section.

## 2. Materials and Methods

### 2.1. Lipids, chemicals and sample preparation

All lipids were purchased in form of powder from Avanti Polar Lipids (Alabaster, AL) and used without further purification. For overview of the chemical structure of the studied glycerophospholipids and sphingolipids, see supplementary Fig. S5. *Cis*-9-tricosene as well as chloroform and methanol (pro analysis grade) were obtained from Sigma-Aldrich (Vienna, Austria). Stock solutions of lipid (10 mg/ml) and tricosene (5 mg/ml) were prepared by dissolving weighted amounts in organic solvent chloroform/methanol (9:1, vol/vol).

Fully hydrated H_II_ phases were prepared as detailed previously [8, 12]. In brief, stock solutions were mixed at appropriate molar ratios and added to 300 *µ*l ultra-pure water (18 MΩ*/*cm^2^, organic solvent/water ratio = 2.55 vol/vol), which was incubated before mixing at 65°C in 20 ml test tubes. The mixture was then quickly mounted onto a modified rapid solvent exchange apparatus [27] to remove the organic solvent within 5 min by adjusting the vaccum pressure to 400 − 500 mbar, using a vortex speed of 600 rpm and an Ar flow of 60 ml/min. The final samples contained 12 wt.% tricosene.

### 2.2. Small-angle X-ray scattering

Small-angle X-ray scattering (SAXS) experiments were performed using a SAXS-pace system (Anton Paar, Graz, Austria), equipped with a 30 W-Genix 3D microfocus X-ray generator (Xenocs, Sassenage, France) with a Cu-anode and an Eiger R 1 M detector system (Dectris, Baden-Daettwil, Switzerland). Alternatively, a SAXSpoint camera (Anton Paar) connected to a MetalJet (Excillum, Kista, Sweden) with a liquid, Ga-rich alloy, jet anode was used. This system too was equipped with an Eiger R1 M detector. Samples were contained in paste cells (Anton Paar) and exposed to X-rays for 30 minutes (6 frames à 5 min) for the SAXSpace, and 2 minutes (12 frames à 10 s) when using the SAXSpoint camera. All samples were equilibrated for 10 minutes at each temperature (TC 150, Anton Paar) prior to measurement starting at the highest measured temperature. This protocol was found to aid the homogeneous distribution of guest lipids in the host matrix. The sample-to-detector distances were 308 mm (SAXSpace) and 700 mm (SAXSpoint). Data were background corrected and integrated using SAXSanalyis (Anton Paar).

### 2.3. Dilatometry

The molecular volume of pure tricosene was determined as a function of temperature using a DMA 5000 M (Anton Paar), which applies the vibrating tube principle for density measurements [28].

### 2.4. Molecular Dynamics Simulations

We performed all-atom molecular dynamics simulations of 128 lipid molecules together with 40 tricosene chains and *N* water molecules. A fraction of *x* of the lipid molecules consisted of DPhPC molecules, thus 1 − *x* accounts for the fraction of DOPE. The lattice constant was fixed to the experimental value, and the pressure was set to 1 atm using a semi-isotropic barostat. The number of water molecules in the system was determined by equating the chemical potential of a water molecule in the simulation system to its value in bulk water. A snapshot of a simulation with *x* = 0, using the *z*-direction as the direction of view, is shown in Fig. 1C. The boundary conditions are periodic in three directions, using a triclinic primary box with an angle of *π/*3 to place the periodic images on a hexagonal grid. We used the CHARMM-36 force field for the DOPE, the DPhPC and the tricosene [29], and TIP3P for the water [30]. The simulations were performed using GROMACS 2019 [31]. Simulation details are discussed in the supplementary information (SI).

**Figure 1:**
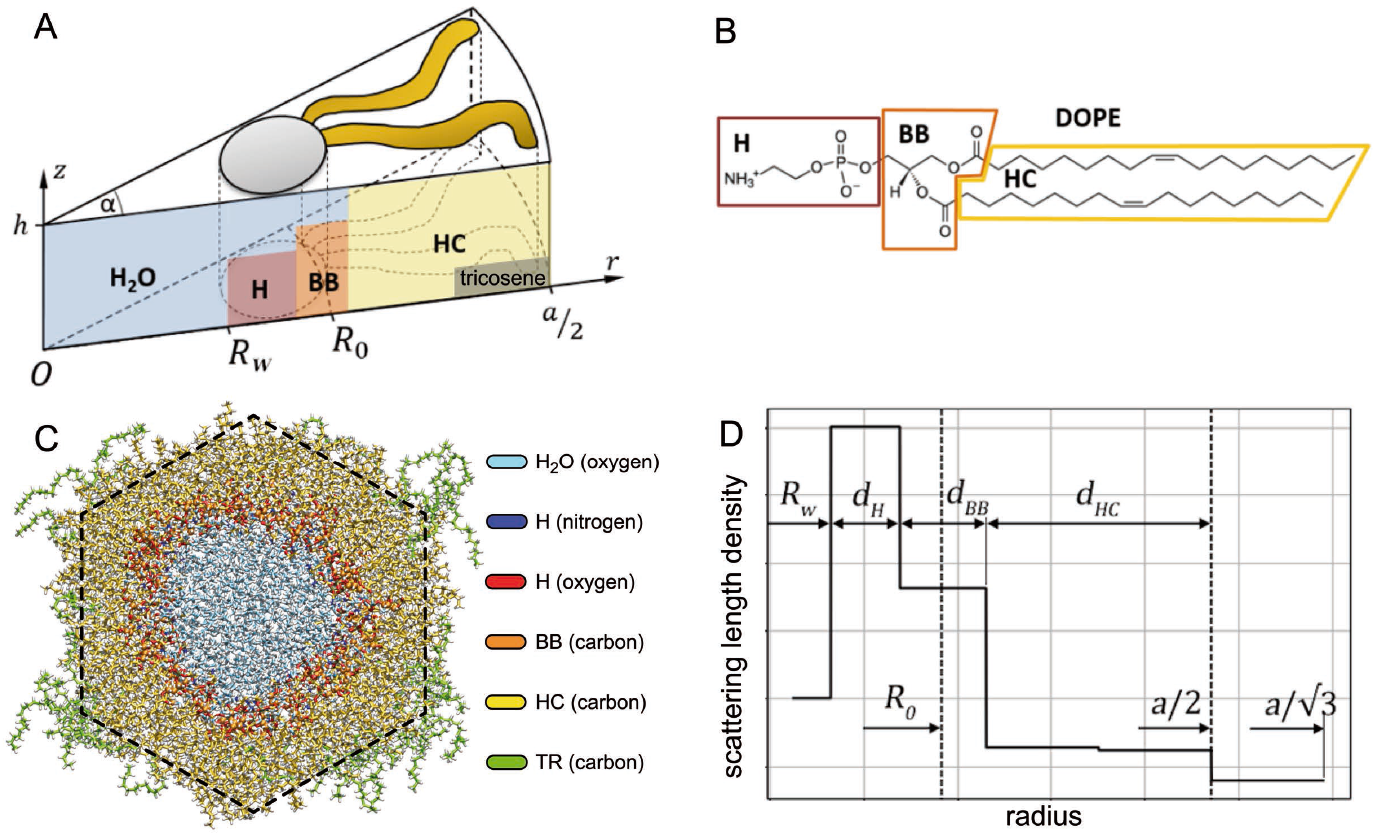
Composition-specific modelling of lipid structure in H_II_ phases. Panel A shows the wedge-like lipid unit cell of radius *a/*2, where *a* is the lattice parameter and *R*_*W*_ is the radius of the water core. Panel B gives an example for the parsing of a lipid molecule. (C) MD simulation snapshot of DOPE (colors: water light blue; headgroup nitrogen: dark blue; headgroup oxygen: red; backbone carbon; orange; hydrocarbon carbon: yellow; tricosene carbon: green). (D) Scheme of the scattering length density profile along the radial axis. *R*_*W*_ is the radius of the water core, *d*_H_ is the thickness of the headgroup layer, *d*_BB_ is the thickness of the backbone layer, and *d*_*HC*_ is the thickness of the hydrocarbon chain layer.

### 3. Theoretical Considerations and Data Analysis

#### 3.1. Non-linear effects on intrinsic curvature from lipid shape

To derive the intrinsic curvature of lipid mixtures forming the H_II_ phase, we consider two wedge-shaped lipid components which are characterized by wedge angles *ω*_*h*_ and *ω*_*g*_ and headgroup areas *A*_0,*h*_ and *A*_0,*g*_ measured in the neutral surface. The subscripts *h* and *g* refer to the host and guest lipid, respectively (Fig. 2A). In our notation, positive angles *ω*_*h*_ and *ω*_*g*_ correspond to lipids with a negative curvature and *vice versa*. We assume that the average headgroup dimension along the axis of the inverted micellar rods is the same for the host and the guest lipid component whereas their effective sizes in the transverse plane, denoted by *b*_*h*_ and *b*_*g*_ (Fig. 2A), are different.

**Figure 2:**
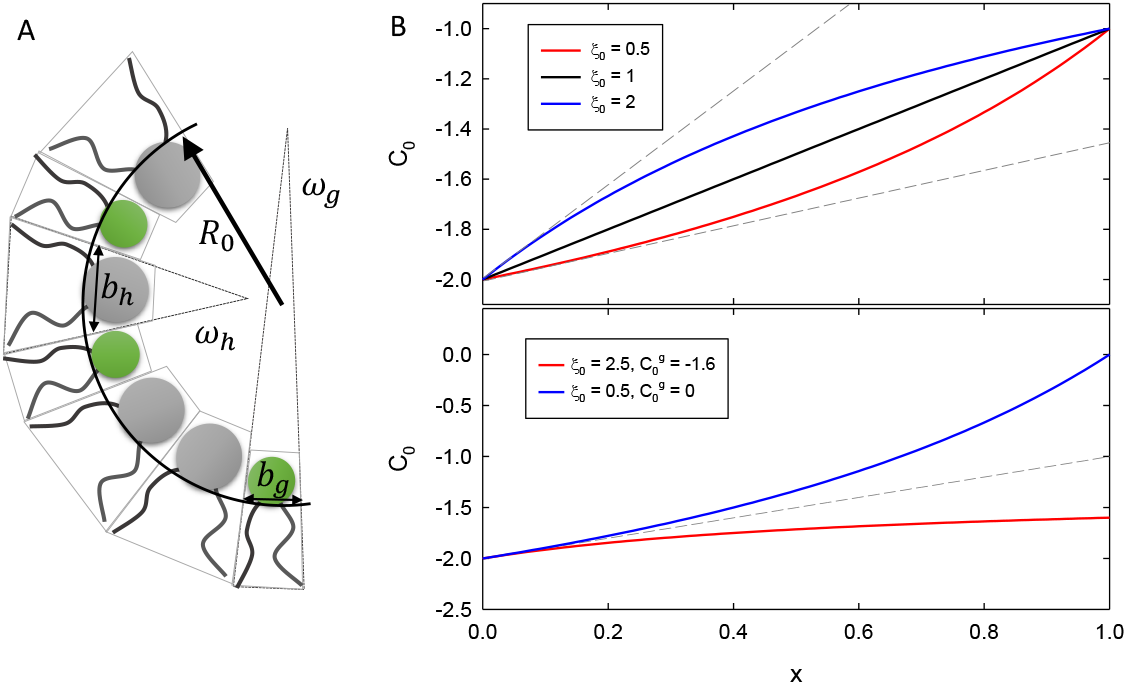
Non-linear curvature mixing. Panel A shows a schematic of the transverse crosssection of a H_II_ phase consisting of two types of wedge-shaped lipids of angles *ω*_*h*_ and *ω*_*g*_, respectively, and effective in-plane headgroup dimensions *b*_*h*_ and *b*_*g*_, respectively. The circular arc represents the neutral surface of radius *R*_0_. For simplicity, both components are sketched with equal tail lengths. Panel B demonstrates the non-linear mixing effects on the intrinsic lipid curvature based on Eq. (3), using 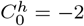 and 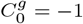 (arbitrary units; top panel) for a few values of *ξ*_0_. Dashed lines indicate linear extrapolations from small *x*. The bottom panel shows two *C*_0_(*x*) profiles that agree at *x ≪* 1 but deviate from each other as *x* is increased.

In terms of the two wedge angles, the total angle subtended by the lipids in the transverse plane is given by

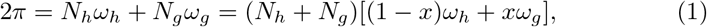

where *N*_*h*_ and *N*_*g*_ are the numbers of host and guest lipids whereas *x* = *N*_*g*_*/*(*N*_*g*_ + *N*_*h*_) is the mole fraction of the guest component. Likewise, the effective perimeter of the transverse-plane cross-section of the neutral surface of the H_II_ phase reads

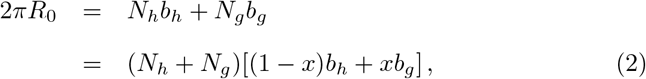

where we assume that *b*_*h*_, *b*_*g*_ ≪ *R*_0_. Now we express *N*_*h*_ + *N*_*g*_ from Eq. (1) to calculate the radius *R*_0_ and from it the intrinsic curvature defined by *C*_0_ = −1*/R*_0_ [32]:

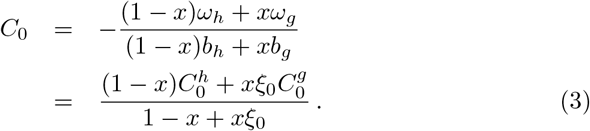

Here 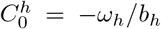 and 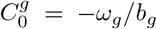 and *ξ*_0_ = *b*_*g*_*/b*_*h*_ is the ratio of the transverse-plane effective sizes of the lipids at the neutral surface. In this expression, *ξ*_0_ leads to a non-linear mixing of the intrinsic curvatures of the two lipids as seen in Fig. 2B. Consequently we refer to *ξ*_0_ as the non-linearity parameter. Note that for *ξ*_0_ = 1, Eq. (3) reduces to the previously used linear formula for the curvature of a mixture 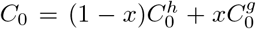 (see, e.g. [14, 16, 17]). Figure 2B demonstrates that if *ξ*_0_ is either considerably smaller or considerably larger than unity, linear extrapolations from low concentrations of the guest lipids lead to a significant over-or underestimate of 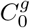, respectively.

### 3.2. Deriving intrinsic lipid curvatures from SAXS

Finding the experimental 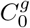 -values using Eq. (3) essentially boils down to finding the neutral plane of a given lipid mixture forming the H_II_ phase. Analogously to our previous reports [17, 12] and in line with MD simulations [33], we assume that *R*_0_ coincides with the position of the lipid backbone. Previously, we also demonstrated how to derive *C*_0_ from a global SAXS data analysis of single lipid species forming a H_II_ phase [12]. In the present work, we generalize this technique to two-component lipid mixtures. Full details of the analysis are given in SI. Here we provide only the most important features of the model.

In brief, the structure of a hexagonal prism can be described using a wedge-shaped lipid unit cell of opening angle *α* and height *h* parsed into four layers, three of which contain quasi-molecular lipid fragments: Headgroup (H), backbone (BB) and hydrocarbons (HC) (Fig. 1A); see Fig. 1B for the parsing of DOPE. Previously, we assumed that tricosene is fully constrained to the interstitial space between the hexagonally packed micellar rods. Interestingly, however, our MD simulations showed that tricosence is able to penetrate also significantly into the HC layer (Fig. 1C). In order to account for this effect, we therefore decided to include tricosene in the terminal half of the HC layer (Fig. 1D); see SI for details.

The intrinsic lipid curvature of the guest molecules is most conveniently derived at the boundary to the HC slab. Assuming that the ratio of transverse-plane lipid dimension is defined through the guest and host lipid areas (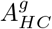 and 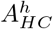) at the hydrocarbon interface, 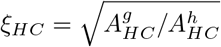, leads to

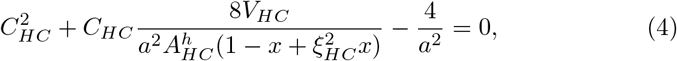

which allows to derive a single solution for *C*_*HC*_, imposing the net curvature to be negative in order to form a H_II_ phase. The curvature of the guest lipid 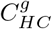 is then derived using Eq. (3), transferred to the HC plane for a known 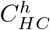. Finally, the intrinsic curvature at the neutral surface is obtained using 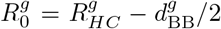. The structural parameters of Eq. (4) are determined by fitting the Fourier transform of the scattering length density profile (Fig. 1D) combined with the hexagonal structure factor to experimental SAXS data of relaxed H_II_ phases using a model that accounts for the full range of scattering vectors *q* (see SI for details).

A specific feature of this model is the need to include a term for the diffuse scattering that does not originate from a pure H_II_ phase. We previously suggested to include scattering from a lipid bilayer [12]. Albeit this term is not of physical interest, it aids in improving the fit of the model to the experimental data. A reevaluation of the global SAXS data analysis for the present work led us to we replace this term by that of a lipid monolayer in case of single component H_II_ phases (see SI). The lipid monolayer can be considered to coat the outer hydrophobic surface of the hexagonally packed inverted micellar bundles and is physically more realistic than a coexisting lamellar phase. Moreover, the quality of fits for DOPE improved significantly (see Sec. 4). However, in case of lipid mixtures, we found that the SAXS data is best described by also including a lipid bilayer. This can be understood by considering that not all guest lipids may partition into the H_II_ phase.

### 3.3. Data fitting

Like in our previous report [12], we applied a Bayesian probability theory for optimization of the adjustable parameters (see SI for details). Within this framework any information on the adjustable parameters is represented in probability distribution functions and the model is formulated in terms of a likelihood function, which is integrated using a Markov Chain Monte Carlo (MCMC) algorithm [34]. In general, parameter distribution functions were drawn from flat prior (starting) distributions. However, for two parameters a better convergence was achieved using Gaussian prior distributions. In particular, the actual guest lipid concentration within the H_II_ phase may differ slightly from its nominal value due to the presence of a coexisting lamellar phase or uncertainties in sample preparation. The corresponding Gaussian prior distributions were therefore centered at the nominal concentrations with a 10% standard deviation. In case of ceramides, we determined *ξ*_*HC*_ from fits to mixtures of DOPE and C16:0 ceramide and used the results as Gaussian prior distribution for all other ceramides. Externally supplied parameters included *V*_H_ and *V*_BB_, which were taken from previous high-resolution studies on lamellar phases formed by phosphatidyl-ethanolamines [35], phosphatidylcholines [36] and sphingomyelin [37]; see Tables S1, S2, and S3 for a complete list of adjustable and supplied parameters.

For illustration, Fig. 3A gives the resulting probability density distribution of *ξ*_*HC*_ for a DOPE:DOPC mixture and demonstrates how the corresponding parameter values and uncertainties are determined. Further, the MCMC analysis allows to screen for parameter correlations (Fig. 3B). In the presently studied systems no significant correlations were observed, which supports the used parametrization of our model. When studying the system for a series of guest-lipid concentrations of the guest lipids, it is further possible to increase the reliability of the obtained structural results by jointly analyzing the data at all concentrations. In particular, *ξ*_*HC*_ and 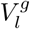 do not depend on the concentration of the guest lipids and thus can be optimized simultaneously for the complete concentration series. In case of sphingolipids, higher stability was achieved by using fixed volumes for the guest lipids in the hexagonal and lamellar phases (Tables S3 and S4). Finally, the derived parameters such as *C*_*HC*_ were then calculated by averaging over all corresponding MCMC trajectories.

**Figure 3:**
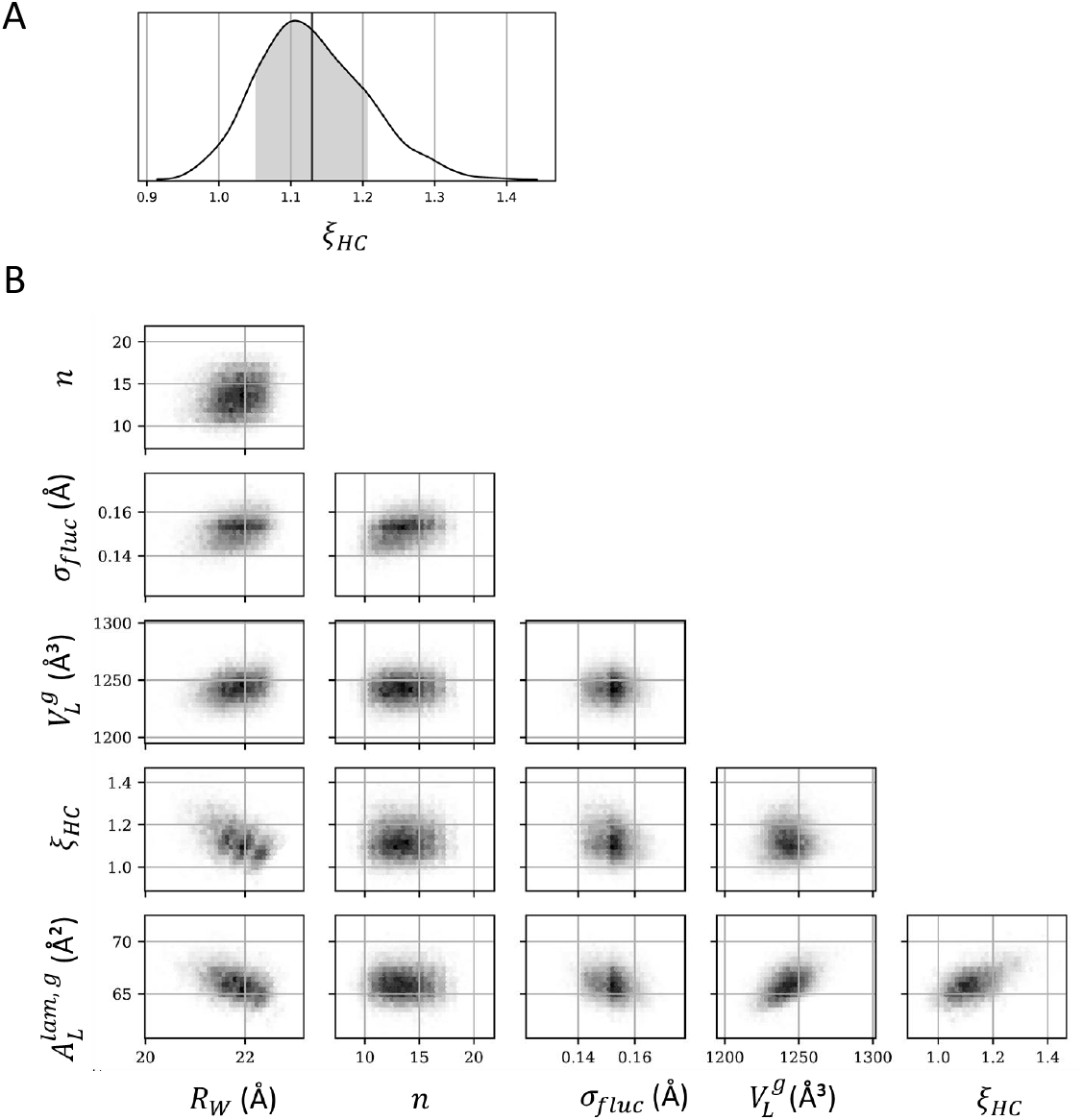
Marginal posterior distributions of DOPC (10 mol%) in H_II_ templates of DOPE (*T* = 35°C). (A): Probability density of the transverse-plane lipid size ratio at the hydrocarbon interface. The gray area in the posterior distribution gives the confidence interval and the solid vertical line is the average value of *ξ*_*HC*_. (B): Pairwise correlations between the average number of hexagonal cells *n*, positional fluctuation of the lipid unit cell σ_*fl*_, volume of the guest lipid 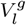, transverse-plane lipid size ratio *ξ*_*HC*_, area of the guest lipid in the lamellar phase 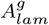, and radius of the water core *R*_*W*_.

### 3.4. Deriving the intrinsic lipid curvature from the simulations

We calculated the electron density of the lipid backbone as defined in Fig. 1B, including both lipid types, as a function of the distance to the central axis of the simulation box, weighting each atom by its atomic number. Fitting a Gaussian function to this density distribution gives the position *R*_0_ and the corresponding curvature. Standard deviations were derived from the fluctuations of the fitted *R*_0_ over time.

## 4. Results

### 4.1. Reevaluation of host lipid structures

The applied changes of the global H_II_ model, i.e. the substitution of the bilayer by a monolayer diffuse scattering contribution, necessitate a check of the most important structural parameters of DOPE and 16:1 PE. Figure 4 shows the corresponding fits for both systems; the results for the structural parameters are reported in Tables S6 and S7. Unlike our previous model [12], the present model faithfully reproduces the near extinction of the (2,1)-peak. The diffuse scattering between the (1,0) and (1,1) reflections is slightly less well accounted for. However, since this contribution does not originate directly from the H_II_ phase, this issue is of less significance. In case of 16:1PE we also added a bilayer contribution, which indeed improved the quality of the fit in this *q*-range. Interestingly, and despite the improved agreement between fit and data the obtained structural parameters for the H_II_ phase are within the experimental error equal to those of our previous report [12]; in particular, 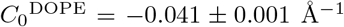 and 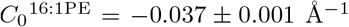 at 35°C. This provides evidence for the robustness of the H_II_ model. Moreover, our MD simulations yielded 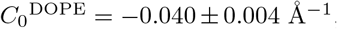, in excellent agreement with the experimental value. The temperature dependence of selected parameters for DOPE is presented in Fig. S6.

**Figure 4:**
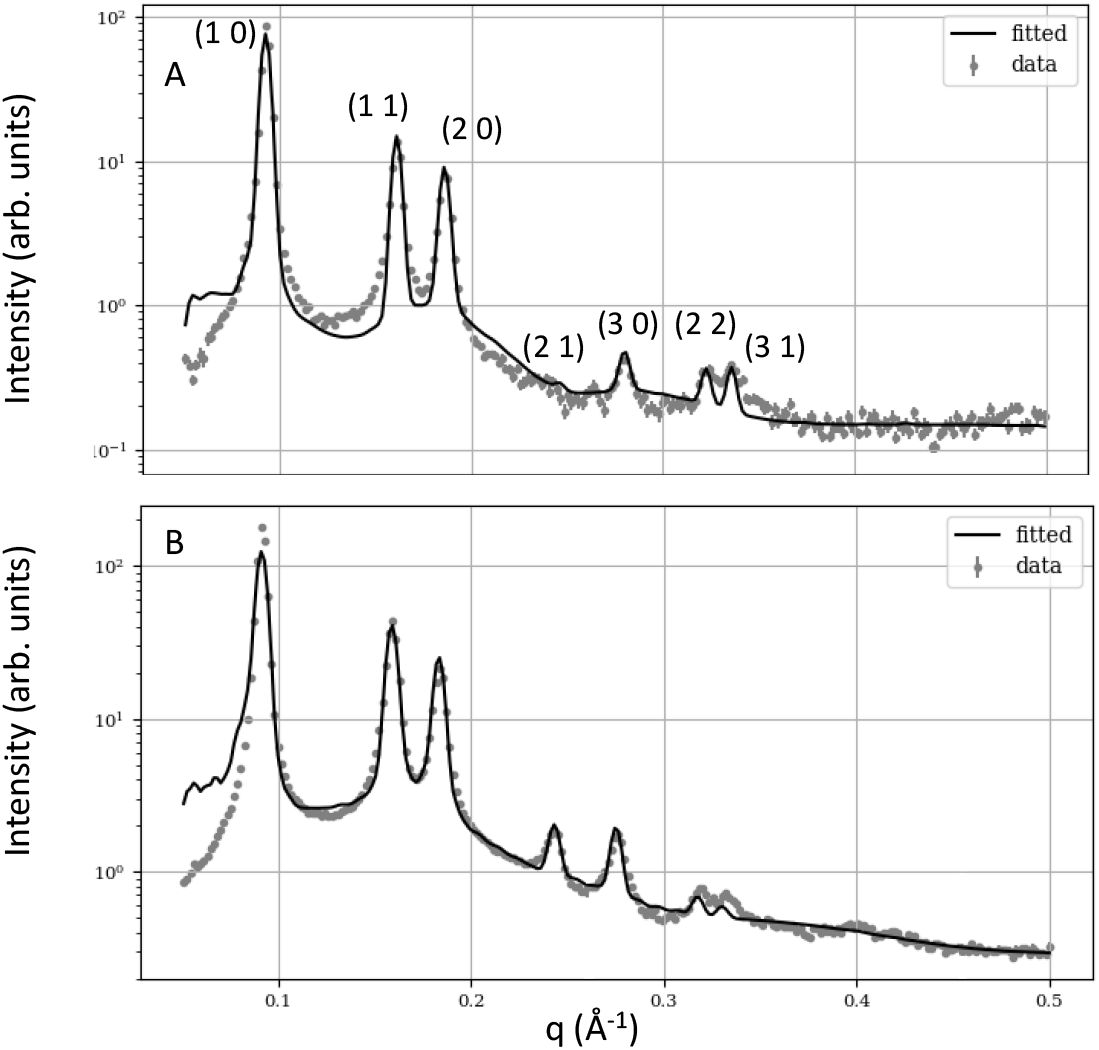
Global analysis of DOPE (A) and 16:1PE (B) at 35°C.

### 4.2. Phosphatidylcholines

We first applied our analysis to phosphatidylcholines with varying hydrocarbon chain composition starting with DPhPC, which should exhibit a significant negative *C*_0_ as well due to its branched hydrocarbons. Figure 5A presents the global fit of DOPE containing 10 mol% DPhPC, which, except for very low *q*-values and the (2,2) and (3,1) reflections in the range *q* = 0.31 − 0.35 Å^−1^, perfectly accounts for the measured intensities. Table S8 shows the corresponding structural parameters of the mixture; for the intrinsic curvature of DPhPC we find 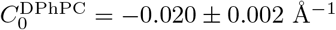.

**Figure 5:**
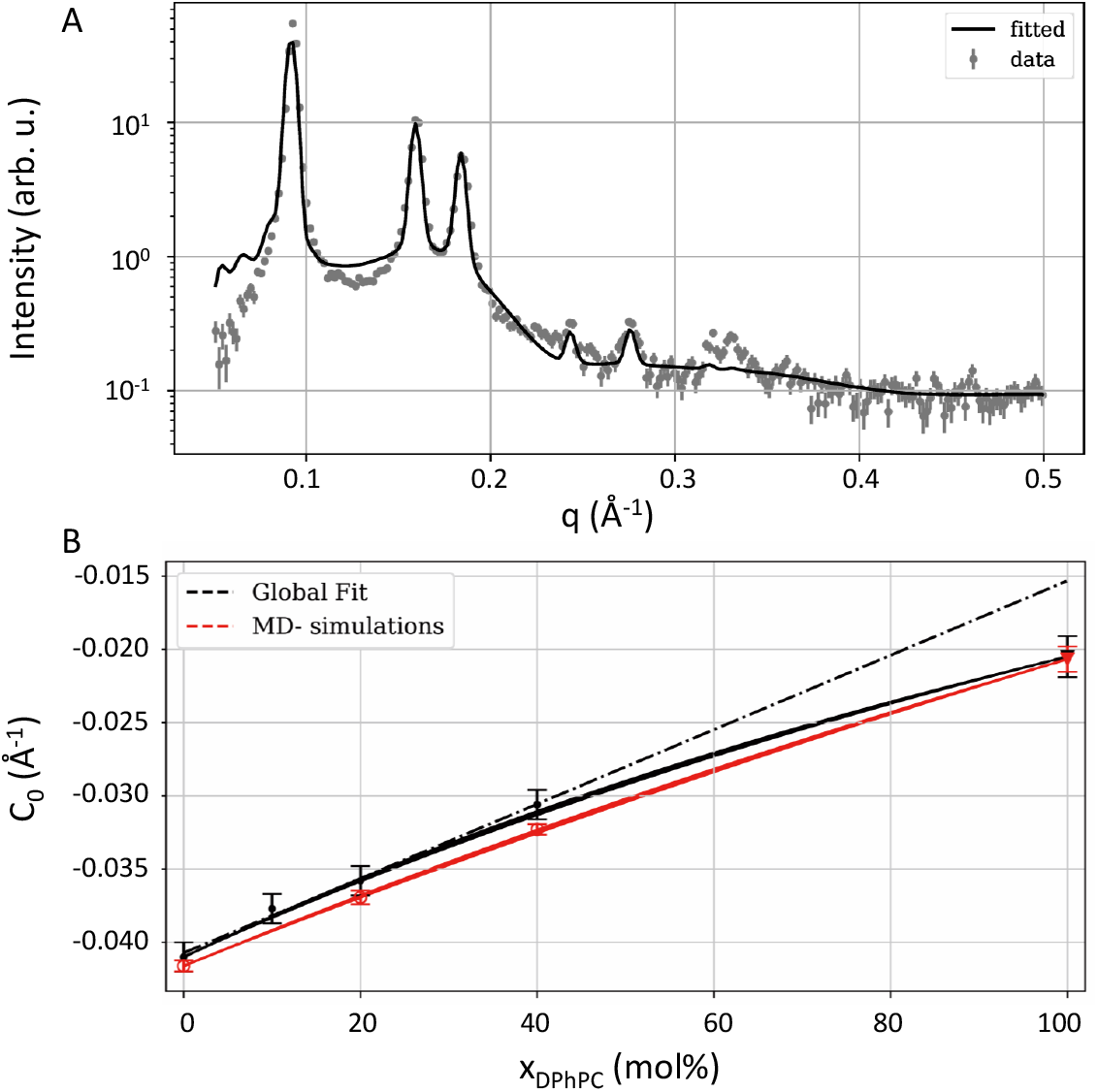
Global analysis of DPhPC guest lipids (10 mol%) at 35°C (panel A). Panel B shows the concentration dependence of the total intrinsic curvature of DOPE/DPhPC mixtures as obtained from the global analysis (black line and symbols) and MD simulations (red line and symbols). The dash-dotted lines shows a linear extrapolation of experimental *C*_0_ data.

We also studied higher concentrations of DPhPC (20 mol% and 40 mol%). These data were jointly analyzed as detailed in Sec. 3.3, adjusting *V*_*L*_ and *ξ*_*HC*_ simultaneously for both data sets. The results of this analysis are shown in Figs. S7A and B as well as in Table S8. Interestingly, 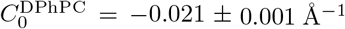 is within the experimental uncertainty equal to the intrinsic curvature obtained by analyzing the 10 mol% sample only, demonstrating the robustness of the model at low guest lipid concentrations. If we extrapolate linearly to 100% DPhPC (i.e. *ξ*_0_ = 1), we find 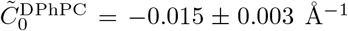 instead, which is significantly shifted toward positive values compared to the non-linear result (Fig. 5B).

DPhPC is also a good candidate to perform a cross-check with MD simulations (Fig. 6). The electron densities retrieved from the simulations show a larger radius of the water column at *x* = 40% (panel B) than at 0% (panel A), which is due to increase of water molecules in equilibrium with increasing DPhPC content. DPhPC has a larger head group than DOPE, and its head groups are located closer to the center of the box, as can be seen by comparing the red dashed and red solid lines in Fig. 6B. The position of the backbone (orange solid and dashed lines) shifts to higher radii with increasing DPhPC content, but shows identical behavior for DPhPC and DOPE. This verifies our assumption of aligned backbones in SAXS data analysis (see Sec. 3.2). The center of the backbone position, *R*_0_, averaged over both lipid types increased from 24 ± 0.3 Å at *x* = 0% to 31 ± 0.3 Å at 40% (Fig. 6C). The shape of the different lipid molecules can be visualized by first rotating the backbone position of all lipids to the horizontal axis. We then averaged their electron density as a function of *r* − *R*_0_ and the coordinate *w* perpendicular to the radial axis, as indicated Fig. 6D. The resulting average electron densities of DOPE and DPhPC signify the larger head group and the ‘heavier’ tails of DPhPC (Fig. 6E). Yet, the spatial distribution of the tails is similar for the two lipid types, but somewhat more extended for DOPE.

**Figure 6:**
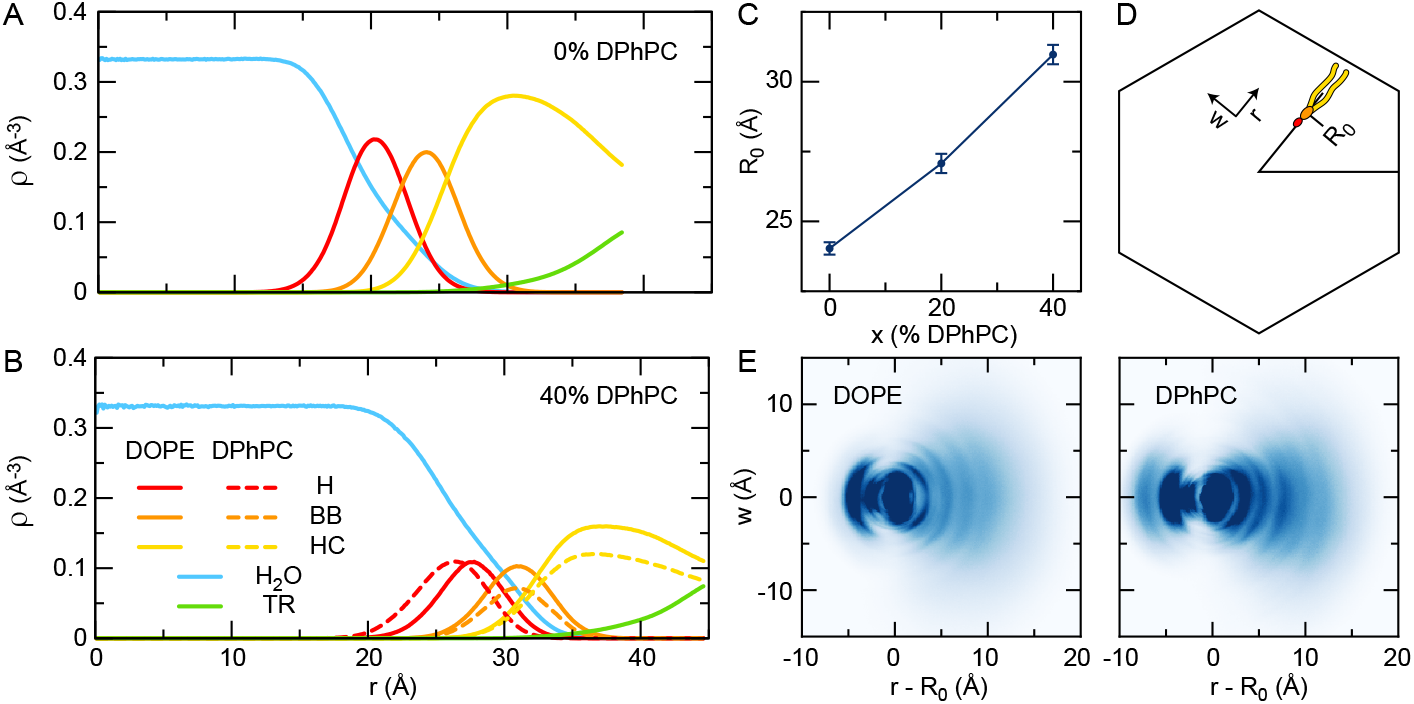
Molecular dynamics simulation results for the electron density *ρ*(*r*) as a function of the radial distance *r* from the center of the water column, split into the contributions from water, lipid head group, backbone, hydrocarbon tails and tricosene, for DOPE (0% DPhPC (A)) and for 40 mol% DPhPC (B). The position *R*_0_ of the backbone as a function of the percentage DPhPC (C). In panel (E), we plot the average electron density of the different molecule types as a function of the radial coordinate *r−R*_0_ and the coordinate *w* perpendicular to the radial direction, as sketched in panel (D).

The area per lipid of DOPE at the neutral plane was calculated from 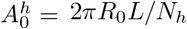, at *x* = 0%, with *L* being the length of the simulation box in axial direction, giving 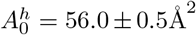, which is lower than our experimental value (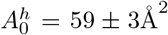; Tab. S6), but still within experimental uncertainty. For the guest lipid, here DPhPC, the area per lipid is calculated from 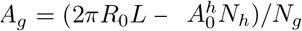, giving *A*_*g*_ = 78.4 ±1.2 at Å ^2^ *x* = 20, and *A*_*g*_ = 76.1 ±1.1 Å ^2^ % at *x* = 40 %. Given the fair agreement between these estimates of *A*_*g*_, we neglect possible non-additive effects for the area per DPhPC molecule. The non-linearity parameter estimated from the MD simulations thus equals *ξ*_0_ = 1.17 ± 0.03. Although this value is lower than the above derived experimental value, it is clearly above 1. The intrinsic curvature of DPhPC, calculated from Eq. (3), equals 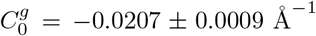 in excellent agreement with our experimental value (see also Fig. 5B). Using *ξ*_0_ = 1 instead, the estimate changes to 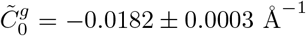, showing that the non-linearity is important to determine the intrinsic curvature of the guest lipids.

Next, we focused on the intrinsic curvatures of DPPC and DOPC. The global H_II_-model agreed well with experimental data at all studied concentrations (Figs. S8 and S9). Again, the obtained 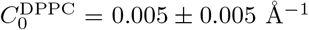 was within the experimental error equal to that resulting from our previous analysis of the same data (Fig. S8D), whereas a linear extrapolation of the globally fitted curvature values gives 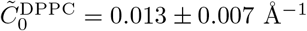. In turn, for DOPC we found 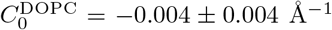. The obtained values of *ξ*_*HC*_ were closer to unity than those for DPhPC (DPPC: 1.12 ± 0.07; DOPC: 1.13 ± 0.08), which is an expression of the fact that DPhPC is more conically shaped than DOPC and DPPC. Previously, we reported smaller intrinsic curvature values, fitting Bragg peak intensities and using linear extrapolations [17].

We then extended the studies to disaturated PCs with shorter (DLPC, DMPC) and longer hydrocarbons (DSPC) as well as to the monounsaturated POPC. Figure 7A shows the variation of *C*_0_ with chain length. The intrinsic curvatures of all phosphatidylcholines with saturated hydrocarbon chains show a weak linear decrease with increasing number of carbon atoms per chain *n*_*C*_: *C*_0_ = −1.15 · 10^−3^Å^−1^ × *n*_*C*_ + 24 · 10^−3^ Å^−1^. Compared to DOPE, the studied phosphatidylcholines exhibited no significant variation of *C*_0_ with temperature (Fig. S10). In case of DMPC, we also used 16:1 PE as H_II_-forming host. The idea was to test whether hydrocarbon chain length mismatch between host and guest lipids affects the curvature measurement. Within experimental uncertainty, however, DMPC guest lipids within 16:1 PE gave the same result (Fig. 7; see also Fig. S11).

**Figure 7:**
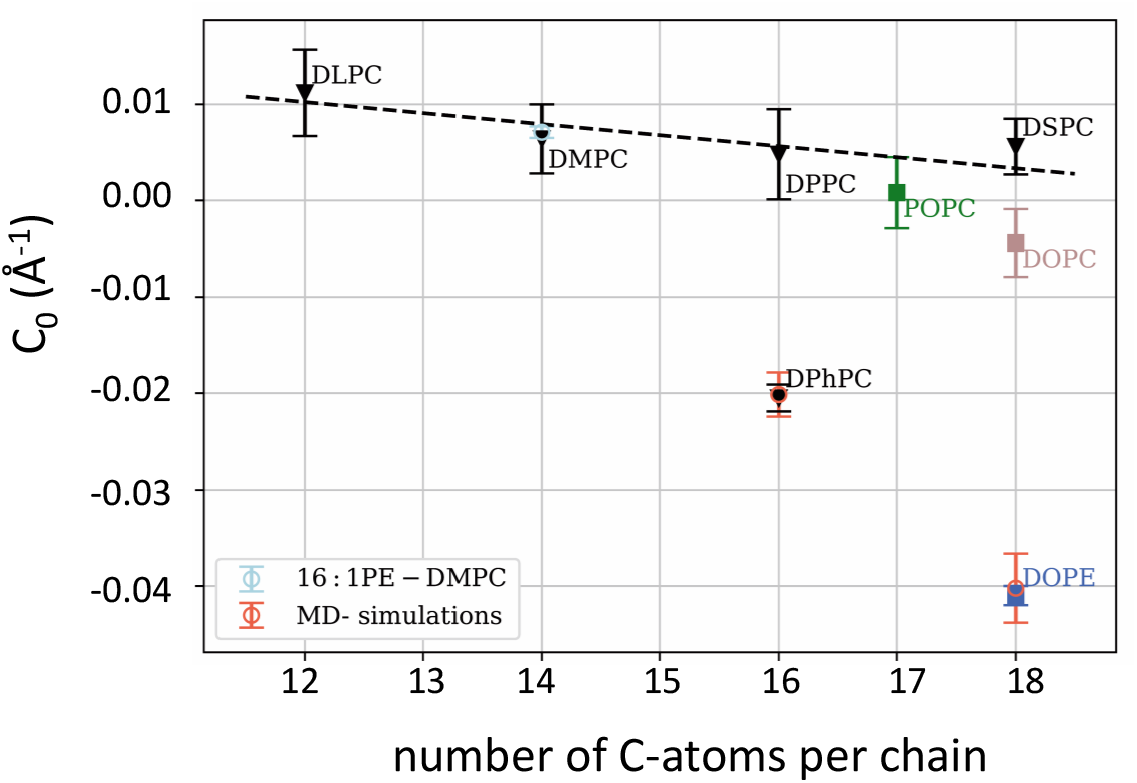
Chain length dependence of intrinsic lipid curvatures of the studied phosphatidylcholines at 35°C.

### 4.3. Sphingomyelin and Ceramides

We next studied the instrinsic lipid curvatures of PSM, as well as C2:0, C6:0, C16:0 and C24:0 ceramide. All sphingolipids are well known for their ability to form extensive intermolecular H-bonding networks, which might lead to non-additive mixing [20]. Out of all presently studied sphingolipids, PSM is deemed to be the best candidate to test for such a scenario, because of its better solubility within lipid aggregates. Indeed, PSM well partitioned into DOPE H_II_ templates up to 20 mol%, while ceramide concentrations were limited to less than 10 mol% as also evidenced in the deviation of the lattice constant from a linear change with guest lipid concentration (Fig. S12).

The combined analysis of all PSM concentrations yielded *ξ*_*HC*_ = 1.38 ± 0.05, similar to that of DPhPC (Table S10). However, the resulting intrinsic lipid curvatures were all slightly positive. Moreover, we observed a dependence of *C*_0_ of PSM on its concentration in DOPE in accordance with non-additive mixing (Fig. 8). In particular, *C*_0_ ∼ 0.012 Å^−1^ for PSM concentrations up to 6.5 mol%, but then dropped by about 50% for 15 mol% PSM and slightly further for 20 mol% PSM, but still remained positive. Interestingly, no significant variation of *C*_0_ with concentration was observed for PSM at 50°C (Fig. S14). Overall, *C*_0_ of PSM did not change with temperature, but remained slightly positive (Fig. 9A).

**Figure 8:**
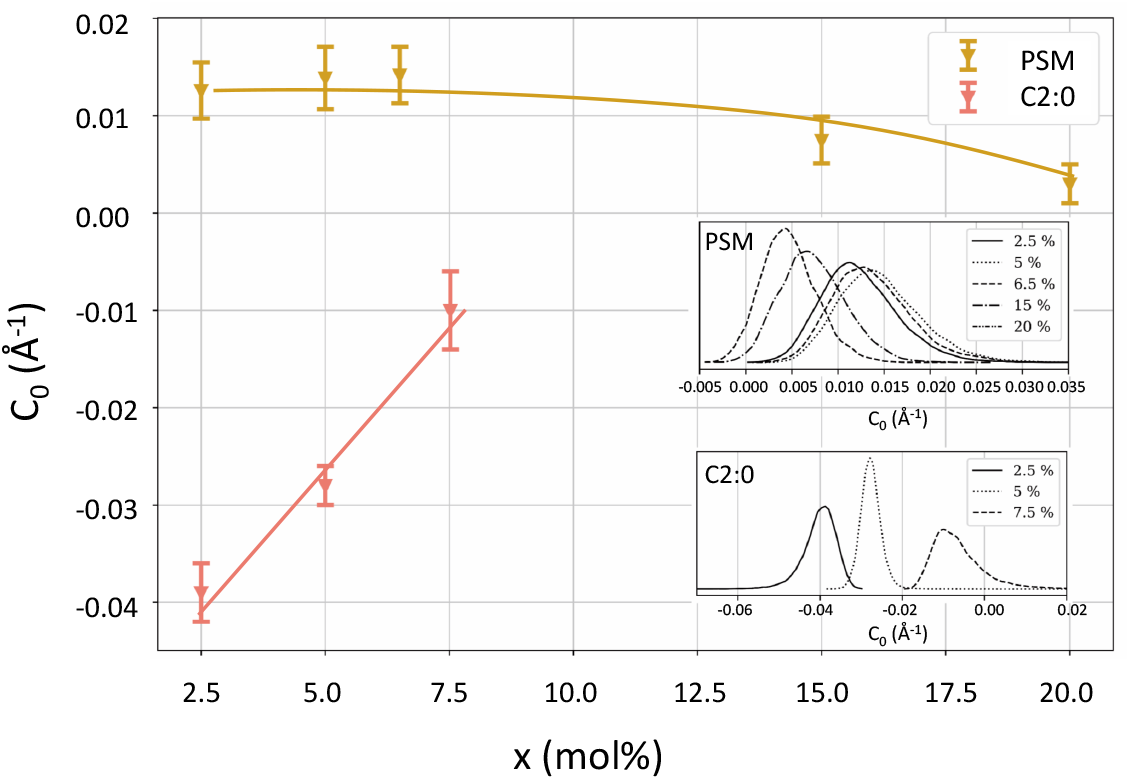
Concentration dependence of intrinsic lipid curvatures of PSM and C2:0 Cer at 35°C. The insets show the corresponding *C*_0_ contributions obtained from a global analysis of SAXS data. Solid lines are a guide to the eye.

**Figure 9:**
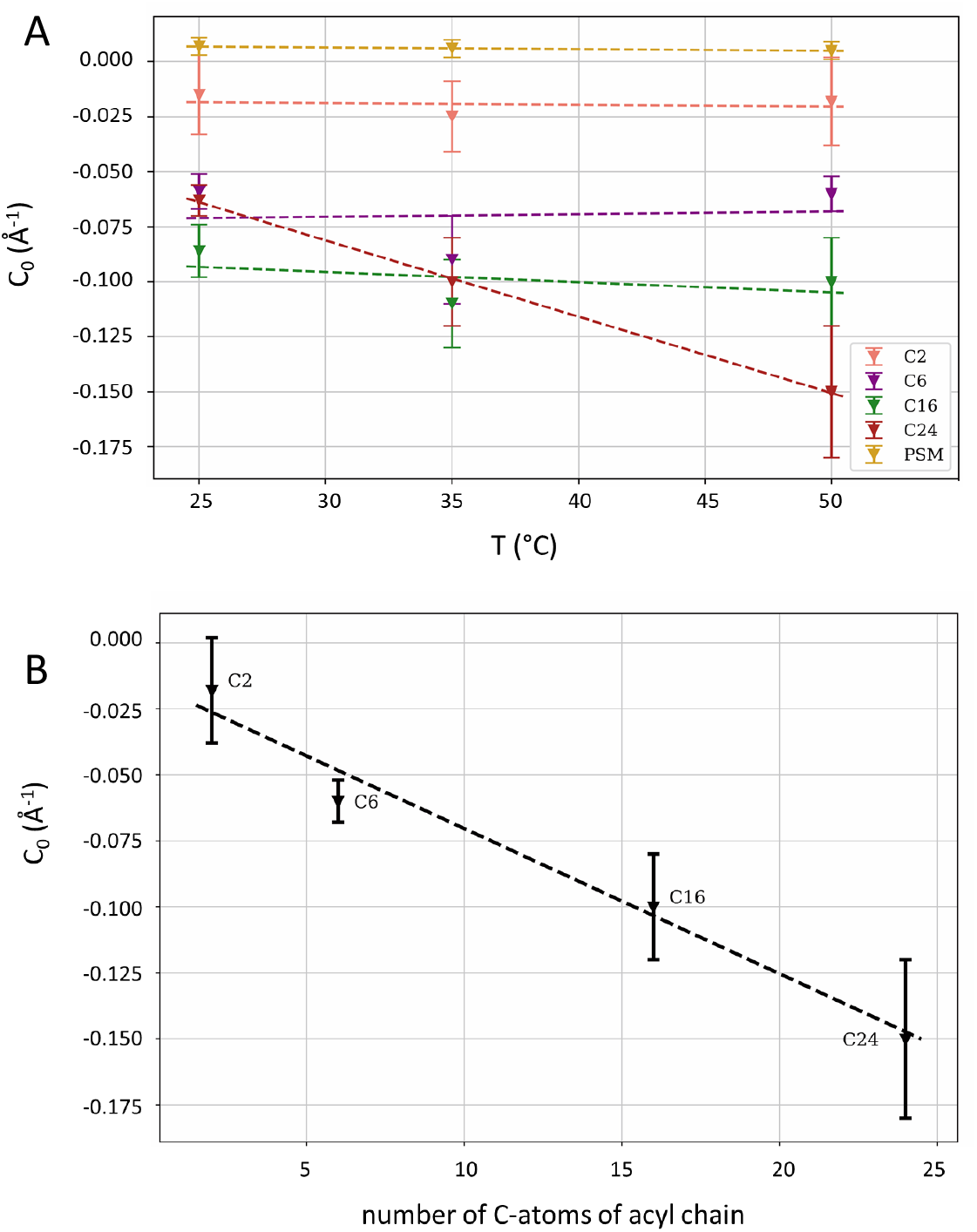
Overview of *C*_0_-values of the studied sphingolipids. Panel A shows the dependence of intrinsic lipid curvatures on the temperature. Average *C*_0_-values are used for PSM and C2:0 Cer. Panel B focuses on ceramides at different lengths of the acyl chain at 50°C.

Ceramides behaved very differently as compared to PSM. Although, we were able to fabricate neat H_II_-phases at least at low ceramide content in DOPE, changes of the lattice constant with ceramide concentration (positive slope for C2:0 Cer; and negative slopes for all other studied Cer; Fig. S12) already indicate significantly different *C*_0_-values. The global analysis of the corresponding H_II_-phase SAXS patterns was challenged by the low solubility of ceramides in DOPE, as well as the low scattering density contrast due to the missing electron rich phosphate group (Fig. S5). Our strategy was therefore to adjust *ξ*_*HC*_ freely in a joint fit of all C16:0 ceramide concentrations and then use this result as prior as detailed in the materials and methods section. Fits and resulting parameters for 50°C are summarized in Figs. S15–S18 and Tabs. S11–S14.

C2:0 ceramide exhibited the lowest ratio of lipid dimension, *ξ*_*HC*_ = 0.3 ± 0.1, which is a result of the small polar headgroup and the short acyl chain. For all other studied ceramides *ξ*_*HC*_ was about two times larger. Intriguingly, we observed a pronounced change of *C*_0_ with concentration for C2:0 Cer (Fig. 8), shifting from *C*_0_ ∼ −0.04 Å^−1^ at 2.5 mol% to *C*_0_ ∼ −0.01 Å^−1^ at 7.5 mol% and 35°C. Except for C24:0 Cer, we observed no concentration dependence for *C*_0_ for all other ceramides (Figs. S16 – S18). C24:0 ceramide was also the only sphingolipid which exhibited a marked temperature dependence for *C*_0_ (Fig. 9 A). Both effects might result from a non-homogenous distribution of C24:0 ceramide in DOPE at low temperatures (see Discussion). In general, increasing the length of the acyl chain of ceramides led to a decrease of *C*_0_ (Fig. 9B).

## 5. Discussion

The global SAXS analysis of H_II_-forming lipid mixtures with DOPE as host developed here advances the determination of intrinsic lipid curvatures of membrane lipids, which normally do not form H_II_ phases. In particular, we extended a previously reported combined compositional data modelling/Bayesian statistical analysis [12] with shape-based packing arguments to address potential non-linear mixing contributions.

Molecular weighted averaging (i.e. linear mixing) is a well-accepted and highly successful concept for estimating the collective properties of planar lipid membranes (e.g. membrane thickness, area per lipid, bending rigidities), see, e.g. [9]. However, this concept cannot be readily transferred to curved geometries (Fig. 2). Indeed, we observed non-linear mixing contributions for all presently studied lipids, albeit least expressed for phosphatidylcholines with saturated hydrocarbons (see e.g. Table S9). Another advantage of the technique developed technique here is the ability to retrieve reliable *C*_0_ values already at low guest lipid concentrations, which may become valuable either if the solubility of the lipid of interest in DOPE is low, or if only little material is available. Here, we applied the technique to a series of phosphatidylcholines and sphingolipids; see Table 1 for a summary of intrinsic lipid curvatures of all studied lipids. From all studied PCs, DPhPC had the most pronounced non-linear effect on mixing with DOPE, which is due to the branched hydrocarbon chain structure, yielding a larger in-plane area at the position of the hydrocarbon interface measured here. In turn, C2:0 ceramide due to its short hydrocarbon chain has a smaller in-plane area than DOPE at the hydrocarbon interface leading to *ξ*_*HC*_ *<* 1 (Table S11).

**Table 1:** List of all here derived intrinsic lipid curvatures at 35°C. Numbers in brackets give the uncertainties for the last digits.

| lipid | $C_0$ ($\text{\AA}^{-1}$) | lipid | $C_0$ ($\text{\AA}^{-1}$) |
| --- | --- | --- | --- |
| DLPC | 0.011 (5) | PSM<sup>b</sup> | 0.010 (6) |
| DMPC<sup>a</sup> | 0.007 (2) | C2:0 Cer<sup>b</sup> | -0.03 (2) |
| DPPC | 0.005 (5) | C6:0 Cer | -0.09 (2) |
| DSPC | 0.008 (3) | C16:0 Cer | -0.11 (2) |
| DOPC | -0.004 (4) | C24:0 Cer<sup>c</sup> | -0.15 (3) |
| POPC | 0.001 (4) | DOPE | -0.041 (1) |
| DPhPC | -0.021 (1) | 16:1PE | -0.037 (1) |

<sup>a</sup> averaged value from measurements in DOPE and 16:1PE
<sup>b</sup> averaged value
<sup>c</sup> 50°C

In terms of intrinsic lipid curvature, we observed that all PCs with saturated hydrocarbons exhibited positive *C*_0_-values, increasing slightly for shorter chain lengths. This indicates stronger lipid headgroup contributions making their effective molecular shape more inverted cone-like. Introducing double bonds or additional methyl groups in the hydrocarbon chain regime reduced *C*_0_ and shifted it to negative values (Table 1).

For the presently studied sphingolipids our improved data analysis yielded positive intrinsic lipid curvatures for PSM, in agreement with previous MD simulations [33, 38]. These results contrast with our previously reported negative *C*_0_-value for egg sphingomyelin [17]. However, these previous data relied on a much less-advanced analysis (integrating Bragg peaks and assuming linear mixing) and much lower guest lipid concentrations, leading to increased uncertainties. The rapid solvent exchange technique applied here [27, 12], which was originally developed for cholesterol rich lipid mixtures [39], avoids passing through a solid phase during sample preparation. This allows to include higher amounts of guest lipid in DOPE and thus aids the reliability of the derived intrinsic lipid curvature.

We also tested our model on its ability to derive non-additive mixing effects, which were proposed to be mediated by interlipid hydrogen-bonding between PSM (or other lipids such as cholesterol) [20]. Indeed, we observed a variation of *C*_0_ of PSM with concentration in DOPE, and even stronger for C2:0 Cer (Fig. 8). Different than predicted by MD simulations for PSM [20], we observe a slightly less positive intrinsic lipid curvatures lipid content. However, the effect observed here is rather small and is also limited by the achievable PSM concentration range in DOPE. Nevertheless, C2:0 Cer exhibited clear non-additive mixing effects and the expected shift of *C*_0_ toward positive values (Fig. 8). This implies that the way ceramides affect the elastic membrane stress is more complex than previously considered. This is to the best of our knowledge the first experimental evidence for the presence of non-additive lipid curvature mixing.

We observed no clear concentration dependence of all other studied ceramides. Instead all ceramides, including C2:0 Cer in the studied concentration range, displayed negative intrinsic curvature values (Tab. 1). Moreover, *C*_0_ values of C6:0, C16:0 and C24:0 Cer were significantly more negative than that of DOPE. C24:0 Cer, however, behaved much differently. For example, C24:0 Cer was the only lipid studied here (including PCs), which exhibited a significant temperature dependence of *C*_0_ (Fig. 9 A). Moreover, values derived at low temperatures were within experimental uncertainty about equal to those of C16:0 Cer. Further, *C*_0_, unlike those of C6:0 and C16:0 Cer suggested some concentration dependence (Fig. S18).

We suggest that the unusual temperature behavior, as well as the change of *C*_0_ with concentration is a result of the long C24:0 hydrocarbon chain. The chain exceeds the C18:1 length of the host lipid. This will lead to a preferential location of the long hydrocarbon within the interstices between the hexagonally packed micellar rods, thus competing with the partitioning tricosene into this region. The resulting non-homogeneous distribution of C24:0 Cer within the H_II_ phase consequently skews our analysis. Moreover, this effect is expected to be more pronounced at lower temperatures due to reduced molecular diffusion. At 50°C instead, *C*_0_ decreased linearly with ceramide chain length (Fig. 9B),suggesting that C24:0 Cer domain formation within the H_II_ is not as much of a concern under these conditions. We therefore propose that the *C*_0_ value derived at this temperature and for 5 mol% concentration corresponds to the true intrinsic lipid curvature of C24:0 Cer.

Given the short hydrocarbon chain of C2:0 Cer and C6:0 Cer a negative *C*_0_ appears as unexpected. However, due to the small headgroup of these lipids, hydrocarbon chain splay contributes much more to the molecular shape. We previously reported *C*_0_ = −0.031 Å^−1^ for C16:0 Cer, based on changes of the H_II_ lattice constant [26]. The large discrepancy from the value reported here points to the subtleties in data analysis. We also note that the formation of a H_II_ phase was claimed for C10:0 Cer [40]. However, close inspection of the reported experimental data suggest rather the formation of lamellar gel phases, consistent also with MD simulations [41].

## 6. Conclusion

A detailed analysis of SAXS patterns of lipid mixtures with H_II_ templates entails deeper and more reliable insight into intrinsic lipid curvatures. In particular, we demonstrated the need to include non-linear mixing contributions to intrinsic curvatures. The applied molecular shape-based approach provides a viable route for experimental exploration. Importantly, the method presented here allows to deduce these values also at low guest lipid concentrations. This may prove particularly valuable if the lipid of interest is available only at low amounts or does not mix well with the template phase. Yet, the restriction to H_II_ phases also incurs limitations in deriving *C*_0_. In particular, fundamental questions relating to non-additive lipid mixing can only be addressed reliably at high amounts of guest lipids, especially when *C*_0_ gradients are low as in the case of PSM. In this case the application of template phases with other topology is deemed more promising.

In the present study we applied our methodology to a series of phosphatidylcholines and sphingolipids, demonstrating the impacts of headgroup and hydrocarbon chain structure on curvature. In particular the significantly negative *C*_0_ values of ceramides at all chain length and the strikingly pronounced nonadditive mixing effect of C2:0 are a manifestation of their ability to strongly to modulate the stored elastic curvature stress in membranes even beyond current considerations. It is most likely that such abilities pertain to several of the reported physiological effects for ceramides [23].

## Acknowledgement

This work was supported by the Austrian Science Funds (FWF) [grant numbers P32514 and P27083]; we also acknowledge the financial support from the Slovenian Research Agency (research core funding No. P1-0055). We also thank F’
selix Goñi for challenging us with suggesting the determination of intrinsic lipid curvatures of ceramides. Mission accomplished.

