## Supplementary Information for "Intrinsic Lipid Curvatures of Mammalian Plasma Membrane Outer Leaflet Lipids and Ceramides"

#### 1. Scattered intensity of relaxed and unoriented $H_{II}$-phases

Following our previous approach [1] it is convenient to decompose the scattered intensity of the $H_{II}$ phase into a hexagonal lattice, described by a structure factor $S(\vec{q})$, and a base given by a prism filled with a inverted rod-like lipid micelle. The prism is described by the form factor $F(\vec{q})$, where $\vec{q}$ is the scattering vector (Fig. S1). Since the length of the prisms is much longer than their diameter, the scattered intensity can be written as [2]

$$I(\vec{q}) \propto |F(\vec{q})|^2 S(\vec{q}), \tag{S1}$$

where

$$S(q,\theta) = 1 + \frac{1}{N_{hex}(n)} e^{-q^2\Delta} \sum_{j\neq k}^{N_{hex}(n)} J_0(q|\vec{R}_j - \vec{R}_k|\sin\theta) \tag{S2}$$

using cylindrical coordinates. In this notation $\theta$ is the angle between the scattering vector $\vec{q}$ and the long axis of the cylinders, $N_{hex}$ is the total number of lattice points for a hexagonal lattice with $n$ rings, and $\Delta$ accounts for thermal fluctuations (Debye-Waller) of the mean positions $\vec{R}_j$ of the lattice points.

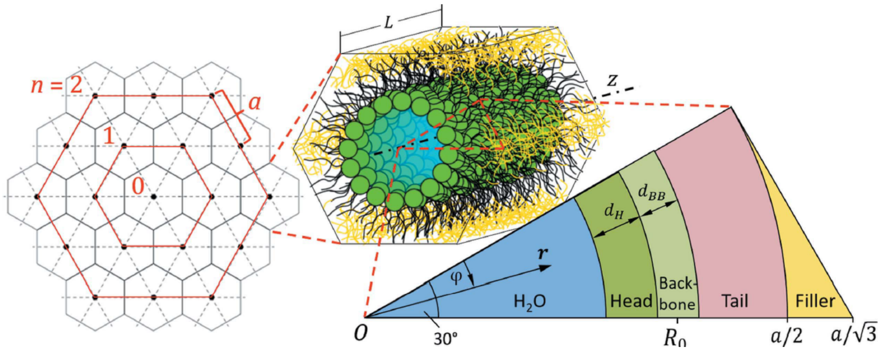


Figure S1: Scheme of the $H_{II}$ phase model. The hexagonal lattice (left side) is defined by its lattice parameter $a$ and the number of rings (lattice order) $n$. Its unit cell, shown in the center, is a regular hexagonal prism of length L and consists of inverted rod-like lipid micelles and a filler molecule (here: tricosene) occupying the interstices. The lipid unit cell is subdivided into areas of different electron densities. $R_0$ denotes the position of the neutral plane. Figure taken from [1] with permission (CC BY 4.0).

The form factor describing a hexagonal prism of length $L$, can be separated into the rod-like inverted lipid micellar part $F_{\text{lipid}}$ and the interstitial part containing tricosene (or other hydrophobic molecules) $F_{\text{inter}}$ [1]

$$\begin{aligned} F(q,\theta) &= f(q,\theta) \int \rho(r,\phi) J_0(q\, r\, \sin\theta) r\, dr\, d\phi \\ &= f(q,\theta) \left[ F_{\text{lipid}}(q,\theta) + F_{\text{inter}}(q,\theta) \right], \end{aligned} \tag{S3}$$

where

$$f(q,\theta) = \frac{4\pi \, \sin\left(\frac{L}{2} q \cos\theta\right)}{q\cos\theta} \tag{S4}$$

and

$$\begin{aligned} F_{\text{lipid}}(q,\theta) &= \int_0^{a/2} r \Delta\rho(r) J_0(q\, r\, \sin\theta)\, dr \\ &= \frac{1}{q\, \sin\theta} \Bigg[ \Delta\rho_M r_M J_1(q\, r_M \sin\theta) + \\ &\qquad + \sum_{k=1}^{M-1} (\Delta\rho_k - \Delta\rho_{k+1} r_k) J_1(q\, r_k\, \sin\theta) \Bigg]. \end{aligned} \tag{S5}$$

Herein $\Delta\rho_k = \rho_k - \rho_W$ is the scattering length density (SLD) relative to water ($\rho_W = 0.33$ Å$^{-3}$) of the $k$'th sector ($k = 1, 2, \ldots M-1, M$), $r_k$ is the radius of the $k$-th cylinder, and $J_0(x)$ and $J_1(x)$ are Bessel functions of the first kind of order 0 and 1, respectively. For the interstices

$$\begin{aligned} F_{\text{inter}}(q,\theta) &= \frac{6}{\pi}\int_{a/2}^{a/\sqrt{3}} dr \int_0^{\pi/6-\arccos(a/2r)} r\Delta\rho_{\text{inter}} J_0(qr\sin\theta)\ d\phi \\ &= \Delta\rho_{\text{inter}}\int_{a/2}^{a/\sqrt{3}}\left[\frac{\pi}{6}-\arccos\left(\frac{a}{2r}\right)\right] rJ_0(qr\sin\theta)\ dr, \end{aligned} \tag{S6}$$

where $\Delta\rho_{\text{inter}} = \rho_{\text{tricosene}} - \rho_W$. This integral needs to be solved numerically. However, $F_{\text{inter}}$ depends only on the lattice constant $a$, which is readily determined from the positions of the Bragg peaks. It suffices to calculate the integral in Eq. (S6) only once for a given $a$-value and store it in an array for all optimization runs.

Fluctuations of the lipid unit cell (Fig. S1) are accounted for by translating the unit cell by distances $x$, which are normally distributed $N(x)$ and centered at $x = 0$ (variance: $\sigma^2_{fluc}$), leading to

$$F_{fluc}(q,\theta) = \int N(x)\, F(q,\theta)\, dx\,. \tag{S7}$$

Finally, the calculation of the total scattering intensity can be substantially simplified by deriving the orientational average, i.e.

$$\begin{aligned} I(q) &= \langle I(q,\theta)\rangle \propto \int_0^{\pi} |F_{fluc}(q,\theta)|^2\, S(q,\theta)\ \sin\theta\, d\theta \\ &\simeq \frac{|F_{fluc}(q)|^2\, S(q)}{q}, \end{aligned} \tag{S8}$$

for which we considered (i) that $f(q,\theta)$ only has a significant contributions at $\theta = \pi/2$ [2] and (ii) that the integral

$$\int_0^{\pi}\left(\frac{2\ \sin(\frac{L}{2}\, q\ \cos\theta)}{q\ \cos\theta}\right)^2 \sin\theta\, d\theta \tag{S9}$$

can be approximated by the Lorentz factor $q^{-1}$. Note that this also leads to a simplified calculation of $F_{\text{lipid}}(q)$ [Eq. (S5)] and $F_{\text{inter}}(q)$ [Eq. (S6)], since all $\sin\theta$-terms become equal to one.

*1.1. Lipid unit cell*

The SLD of each layer $k$ (Figs. **??** and S1) is given by $\rho_k = n_k^e/V_k$, where $n_k^e$ is the total number of electrons of the $k$-th sector for SAXS and

$$V_k = \frac{\hat{A}(r_{k+1}^2 - r_k^2)}{2} \tag{S10}$$

is the theoretical volume of the $k$-th layer, which are filled with the quasi molecular groups according to the applied lipid parsing scheme (here $k$ = H, BB, HC). The mantle area of a sector with unity radius $\hat{A} = h\,\alpha$ is derived from the HC layer using

$$\hat{A} = \frac{2(V_{HC} + V_{tr}^p)}{a^2/4 - R_{HC}^2}, \tag{S11}$$

where $R_{HC}$ is the radius at the boundary of the BB and HC layers and $V_{tr}^p = \Delta\bar{v} \times V_L$ is volume of tricosene, partitioned into the HC sector. Here $V_L$ is the molecular lipid volume and

$$\Delta\bar{v} = \begin{cases} \bar{v}_{\text{tricosene}} - \bar{v}_{\text{inter}}\ , & \text{if } \bar{v}_{\text{tricosene}} > \bar{v}_{\text{inter}} \\ 0\ , & \text{otherwise} \end{cases} \tag{S12}$$

is the volume fraction of tricosene in excess of the available interstitial space, where

$$\bar{v}_{\text{inter}} = \frac{2\sqrt{3} - \pi}{2\sqrt{3} - \frac{4}{a^2}R_w^2\pi}. \tag{S13}$$

is the volume fraction of interstices and

$$\bar{v}_{\text{tricosene}} = \frac{m_{\text{tricosene}}/\rho_{\text{tricosene}}}{m_{\text{lipid}}/\rho_{\text{lipid}} + m_{\text{tricoene}}/\rho_{\text{tricosene}}} \tag{S14}$$

is the total volume fraction of tricosene of the sample, defined by the amount of tricosene used in the preparation (12 mol%). In order to estimate the effect of tricosene on the SLD of the HC sector, we assumed that tricosene partitions primarily in the $HC_2 \equiv [(a - d_{HC})/2; a/2]$ region and we calculate the total volume of this subsector $V_{\text{HC}_2}$ by applying Eq. (S10). This gives the corresponding volume fraction $\hat{v}_{\text{tricosene}} = V_{\text{tricosene}}^p/V_{\text{HC}_2}$ of tricosene, which allows to calculate the modified SLD using

$$\rho_{\text{HC}_2} = \rho_{HC}(1 - \hat{v}_{\text{tricosene}}) + \rho_{\text{tricosene}}\hat{v}_{\text{tricosene}}, \tag{S15}$$

where $\rho_{\text{tricosene}}$ is the SLD of tricosene, which was measured as a function of temperature using dilatometry (Table S5).

In order to calculate the SLDs of the headgroup and backbone sectors we need to consider that both contain also water, where the number of bound water molecules

$$n_W^j = \frac{V_j - V_j^L}{V_W}, \tag{S16}$$

with $j = \text{H, BB}$, where $V_j^L$ refers to the lipid headgroup and backbone volumes, and $V_W = 30$ Å$^3$ is the molecular volume of water. This leads to

$$\rho_j = \frac{n_{j,L}^e + n_W^j n_{H_2O}^e}{V_j}, \tag{S17}$$

where $n_{j,L}^e$ is the number of electrons of the lipid headgroup or backbone, and $n_{\text{H}_2\text{O}}^e = 18$ is the number of electrons of $H_2O$.

Finally, the thicknesses of the individual sectors are given by

$$d_{\text{H}} = R_0 - \frac{d_{\text{BB}}}{2} - R_W \tag{S18}$$

and

$$d_{HC} = \frac{a}{2} - R_0 - \frac{d_{\text{BB}}}{2} \ , \tag{S19}$$

where $R_W$ and $R_0$ are obtained from the global analysis.

For the two-component mixtures, we redefine the SLD by the molecular average of its constituent lipids, i.e., $\rho_k = (1-x)\rho_{k,h} + x\rho_{k,g}$, assuming that lipids align at the HC interface. In Fig. S2 we compare the resulting radial electron density for DOPE with corresponding MD simulations.

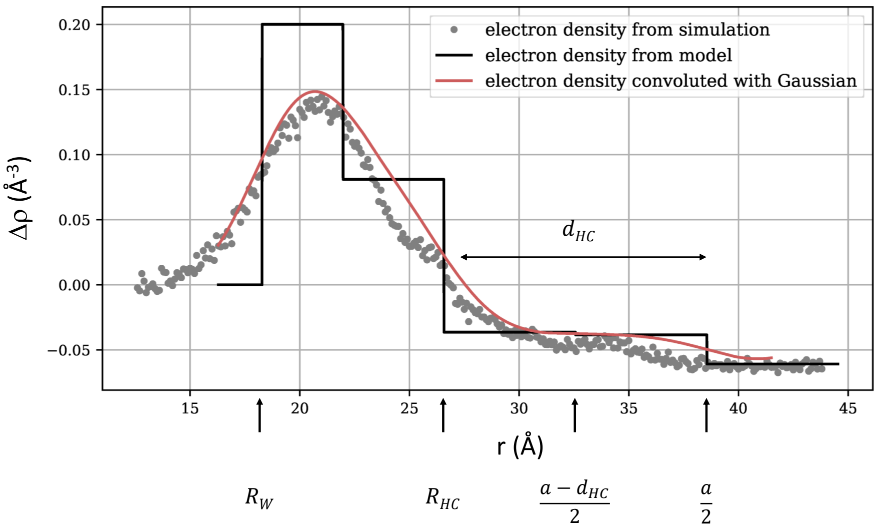


Figure S2: Comparison of the radial SLD slab model for DOPE (35°C) with MD simulations. The red line shows the a convolution of the SLD profile with a Gaussian (width: 20 Å). The small step in the sector model at $(a - d_{HC})/2$ is due to the partitioning of tricosene.

For guest lipids, which differ in their backbone structure from host lipids, such as the here studied sphingolipids, we derived an average backbone width by weighting the backbone widths of the host and guest lipids, $d^h_{\rm BB}$ and $d^g_{\rm BB}$, by their in-plane contributions according to Eq. (**??**). This yields

$$\begin{aligned} \overline{d_{\rm BB}} &= \frac{N_h b_h}{N_h b_h + N_g b_g} d^h_{\rm BB} + \frac{N_g b_g}{N_h b_h + N_g b_g} d^g_{\rm BB} = \\ &= \frac{(1-x) d^h_{\rm BB} + x \xi_0 d^g_{\rm BB}}{(1-x) + x\xi_0}, \end{aligned} \tag{S20}$$

using $\xi_0 = b_b/b_a$.

#### *1.2. Intrinsic curvature of guest lipids*

We first consider again the mantle area of a sector with unity radius, $\hat{A} = h\,\alpha$, which allows us to compute the cross-section area per lipid at any position along the radial axis, e.g. at the neutral surface $A_0 = \hat{A} R_0$. The intrinsic curvature of the guest molecule is most conveniently derived by considering $R_{HC}$. Assuming that the area at $R_{HC}$ is given by the molecular weighted areas of host and guest lipid and $\xi_{HC} = \sqrt{A^g_{HC}/A^h_{HC}}$, we find that

$$\hat{A} = A_{HC}/R_{HC} = -C_{HC}\left(1 - x + \xi^2_{HC}x\right) A^h_{HC}, \tag{S21}$$

which upon combining with Eq. (S11) gives a quadratic equation for the total intrinsic curvature measured at the hydrocarbon interface

$$C^2_{HC} + C_{HC}\frac{8V_{HC}}{a^2 A^h_{HC}\left(1 - x + \xi^2_{HC}x\right)} - \frac{4}{a^2} = 0 \tag{S22}$$

with the discriminant

$$\delta = \left[\frac{8V_{HC}}{a^2 A^h_{HC}\left(1 - x + \xi^2_{HC}x\right)}\right]^2 - \frac{16}{a^2}. \tag{S23}$$

Only $\delta > 0$ gives physically realistic solutions. Moreover, from the two possible solutions of Eq. (S22) we accept only the one which gives an inverted micellar structure, i.e. with a net negative curvature

$$C_{HC} = -\frac{4V_{HC}}{a^2 A^h_{HC}\left(1 - x + \xi^2_{HC}x\right)} - \frac{\sqrt{\delta}}{2}. \tag{S24}$$

Transferring Eq. (**??**) to the HC plane then allows to calculate $C^g_{HC}$ for a known $C^h_{HC}$. All solutions are readily projected onto the neutral surface using $R_0 = R_{HC} - d_{\mathrm{BB}}/2$.

### 2. Modelling diffuse scattering contributions

The observed scattered intensity is not fully described by Eq. (S2). Previously we used a bilayer form factor to account for the additional diffuse scattering [1]. Such a contribution is difficult to justify in the absence of lipid mixtures (e.g. DOPE alone). In Ref. [1] we considered for single phospholipid systems a monolayer coating the outer $\mathrm{H_{II}}$ layer to reduce the energetic penalty of exposing the hydrocarbons to the excess water phase. Using a simple two-slab model accounting for the electron density of the lipid headgroup $\rho^{\mathrm{lam}}_H$ and hydrocarbon region $\rho^{\mathrm{lam}}_{HC}$, the corresponding form factor

$$\begin{aligned} F_{\mathrm{mono}}(q) &= 4\pi^2 \int_{-\infty}^{\infty} \Delta\rho_{\mathrm{lam}}(z)\, \mathrm{e}^{iqz}\, dz \\ &= \frac{4\pi^2 i}{q}\left[\Delta\rho^{\mathrm{lam}}_{HC}\left(\mathrm{e}^{-iqd^{\mathrm{lam}}_{HC}} - 1\right) + \Delta\rho^{\mathrm{lam}}_{H}\left(1 - \mathrm{e}^{iqd^{\mathrm{lam}}_{\mathrm{H}}}\right)\right], \end{aligned} \tag{S25}$$

where $\rho_{\mathrm{H}}^{\mathrm{lam}}$ and $\rho_{HC}^{\mathrm{lam}}$ are defined by the lipid volume of each slab, where $V_{H}^{\mathrm{lam}} = V_{\mathrm{H}}$ and $V_{HC}^{\mathrm{lam}} = V_L - V_H^{\mathrm{lam}}$, with $V_L$ being the total lipid volume, which, for reasons of simplicity and over-fitting, was assumed to have the same value in the lamellar and $\mathrm{H}_{\mathrm{II}}$ phases. The thicknesses of the individual slabs can be also estimated from the $\mathrm{H}_{\mathrm{II}}$ using

$$d_{\mathrm{H}}^{\mathrm{lam}} \approx d_{\mathrm{H}} + d_{\mathrm{BB}} \;\text{ and }\; d_{HC}^{\mathrm{lam}} = d_m - d_{\mathrm{H}}^{\mathrm{lam}}, \tag{S26}$$

where

$$d_m \approx \frac{V_L + n_W^{\mathrm{lam}} V_W}{A_L^{\mathrm{lam}}}, \tag{S27}$$

and

$$n_W^{\mathrm{lam}} \approx \frac{d_m A_L^{\mathrm{lam}} - V_H^{\mathrm{lam}}}{V_W} \tag{S28}$$

are the thickness of the lipid monolayer and number of water molecules per lipid, respectively. The area per lipid in the lamellar phase $A_L^{\mathrm{lam}}$ is an adjustable parameter.

In the case of binary phospholipid mixtures, not all guest lipids may partition into the $\mathrm{H}_{\mathrm{II}}$ phase. To account for this scenario we also included a bilayer form factor

$$\begin{aligned} F_{\mathrm{bi}}(q) \;=\; & -\frac{4\pi^2 i}{q}\Big\{\Delta\rho_{HC}^{\mathrm{lam}}\left[\mathrm{e}^{iq\left(d_{\mathrm{H}}^{\mathrm{lam}}+2d_{HC}^{\mathrm{lam}}\right)} - \mathrm{e}^{iqd_{\mathrm{H}}^{\mathrm{lam}}}\right] + \\ & + \Delta\rho_{\mathrm{H}}^{\mathrm{lam}}\left[e^{iqd_{\mathrm{H}}^{\mathrm{lam}}} - 1 + \mathrm{e}^{i2q\left(d_{\mathrm{H}}^{lam}+d_{HC}^{\mathrm{lam}}\right)} - \mathrm{e}^{iq\left(d_{\mathrm{H}}^{\mathrm{lam}}+2d_{HC}^{\mathrm{lam}}\right)}\right]\Big\}. \end{aligned} \tag{S29}$$

Finally, we assumed that $F_{\mathrm{bi}}$ is entirely determined by the parameters of the guest lipid.

### 3. Full model

The final orientational averaged model for the scattered intensity considers all contributions detailed above, i.e.,

$$\begin{aligned} I(q) &= \Gamma\Big[|F_{fluc}(q)|^2\, S(q) + \\ &+ 2c_{\text{mono}}\, F_{mono}(q)F_{fluc}(q)s(q) + \\ &+ c^2_{\text{mono}}|F_{\text{mono}}(q)|^2 + c^2_{bi}|F_{bi}(q)|^2\Big] + I_{\text{inc}}, \end{aligned} \tag{S30}$$

where the second term considers cross-correlations between the monolayer and the $H_{II}$ phase with

$$s(q,\theta) = \frac{1}{\sqrt{N_{\text{hex}}(n)}} e^{-q^2\Delta/2} \sum_{j}^{N_{\text{hex}}(n)} J_0(q|\vec{R}_j|). \tag{S31}$$

The factors $c_{\text{mono}}$ and $c_{\text{bi}}$ scale the monolayer or bilayer contributions to the scattered intensity. Note that $c_{\text{bi}} = 0$ for single phospholipid component samples. Finally $\Gamma$ is the instrumental scaling constant and $I_{\text{inc}}$ the incoherent scattering contribution. Figure S3 presents the contributions of all structure and form factors to the global scattered intensity of DOPE.

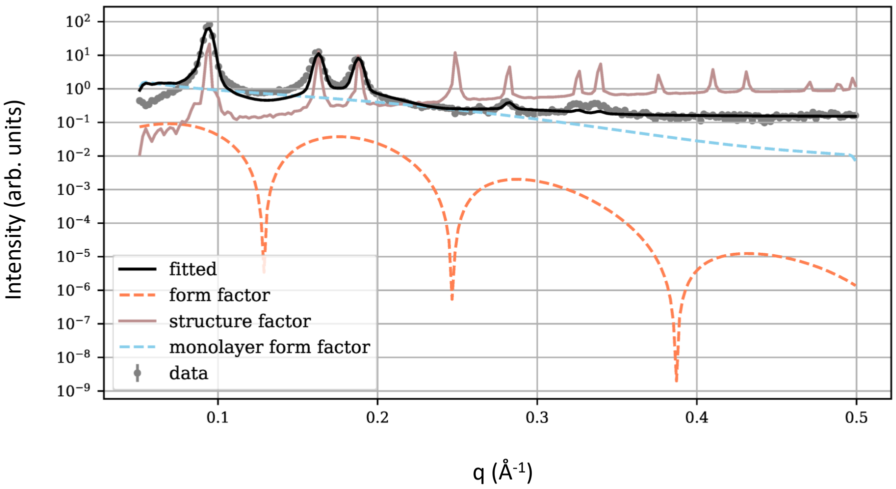


Figure S3: Contributions of $F_{fluc}(q)$ (orange dashed line), $S(q)$ (red line) and $F_{\text{mono}}$ (blue dashed line) to the global $H_{II}$ analysis of DOPE at 35°C (black line).

### 4. Estimating sphingolipid volumes

Increased stability of the algorithm for sphingolipid containing mixtures was achieved by fixing volumes within the $H_{II}$ or lamellar phases (Tables S3, and S4). For ceramides the volume of the polar headgroup was derived by comparing to the lipid volume of PSM [3] and C16:0 Cer [4] at equal temperature. The volumes of all other Cer molecules were calculated starting from the corresponding PSM value by first substituting the headgroup volume and then adding or subtracting the volume of methylene groups ($V_{CH2} = 27$ Å$^3$) [3], according to the given fatty acid chain length.

### 5. Parameter estimation using Bayesian probability theory

Analogously to our previous report, we decided to use Bayesian probability theory, which allows to derive realistic errors for adjustable parameters and yields insight on parameter correlations. Here we just present the most important aspects of this analysis. For details, see Ref. [1] and references therein. In brief, application of Bayes' theorem allows us to derive the probability distribution $p(\hat{\mathbf{a}}|\mathbf{I}, \sigma, \mathcal{I})$ – also called the posterior – of adjustable parameters $\hat{\mathbf{a}}$ of a specific model to fit a given set of experimental data $\mathbf{I}$ with standard deviations $\sigma$ and additional information $\mathcal{I}$.

$$\overbrace{p(\hat{\mathbf{a}}|\mathbf{I}, \sigma, \mathcal{I})}^{\text{posterior}} \propto \overbrace{p(\mathbf{I}|\hat{\mathbf{a}}, \sigma, \mathcal{I})}^{\text{likelihood}} \overbrace{p(\hat{\mathbf{a}}|\mathcal{I})}^{\text{prior}} \ . \tag{S32}$$

The prior probability $p(\hat{\mathbf{a}}|\mathcal{I})$ represents the prior knowledge about $\hat{\mathbf{a}}$, e.g. constraints. The likelihood $p(\mathbf{I}|\hat{\mathbf{a}}, \sigma, \mathcal{I})$ in turn is the probability of measuring $\mathbf{I}$, given $\hat{\mathbf{a}}$ and $\sigma$. Except for the concentration $x$ of the host lipid, all prior probabilities $p(\hat{\mathbf{a}}|\mathcal{I})$ were assumed to be uniformly distributed between lower $\hat{\mathbf{a}}_{\text{min}}$ and upper $\hat{\mathbf{a}}_{\text{max}}$ constraints for all parameters with no interdependence. Due to the presence of a lamellar phase in case of lipid mixtures, the exact guest lipid concentration $x$ in the $H_{II}$ phase is unknown. To account for this uncertainty, we chose a Gaussian prior probability for $x$, centered at the nominal/desired $x$-value with a 10% standard deviation. This probability was, however, not varied during the optimization runs, but was found to keep the fitting procedure more flexible and helped to avoid local minima.

The likelihood [Eq. (S32)] can be connected to the multivariate Gaussian

$$\mathcal{N}(\mathbf{I}|\hat{\mathbf{a}},\sigma) = \prod_{i=1}^{N_q} \frac{1}{\sigma_i\sqrt{2\pi}} \exp\left[-\frac{1}{2\sigma_i^2}(I(q_i|\hat{\mathbf{a}}) - I_i^{\mathrm{obs}})^2\right] \tag{S33}$$

where $I_i^{\mathrm{obs}}$ denotes the observed intensity at $q_i$ and $I_i$, the intensity calculated from the analytical model [Eq. (S30)].

A suitable technique for performing the integrals that connect likelihood and priors to sample the posterior [Eq. (S32)] is the Markov Chain Monte Carlo (MCMC), which is based on constructing a so called Markov chain with the desired distribution of $\hat{\mathbf{a}}$ in equilibrium. We used 'slice sampling' for generating Markov chains [5] and applied the python library 'PyMC3' [6] for parameter optimization. From this analysis we obtained the reported averages of a given adjustable parameter or in general the observable $\mathcal{O}$, which can be a combination of several adjustable parameters by

$$\langle\mathcal{O}(\hat{\mathbf{a}})\rangle \approx \frac{1}{N_{\mathrm{Markov}}} \sum_{k=1}^{N_{\mathrm{Markov}}} \mathcal{O}(\hat{\mathbf{a}}^k)\ , \tag{S34}$$

where $N_{\mathrm{Markov}}$ is the number of uncorrelated Markov Chain elements. The confidence intervals were estimated from

$$\Delta\mathcal{O} := \frac{\sigma_{\mathcal{O}}}{\sqrt{N_{\mathrm{Markov}}}} = \sqrt{\frac{\langle\mathcal{O}(\theta)^2\rangle - \langle\mathcal{O}(\theta)\rangle^2}{N_{\mathrm{Markov}}}}. \tag{S35}$$

To optimize the acceptance during fitting the error derived directly from the experiment was modified with a simple multiplicative scaling factor and a weight

$$\Delta I(q) = \mathrm{e}^{-\gamma(q-q_{\min})}, \tag{S36}$$

where $\gamma$ is also optimized during the MCMC run and $q_{min}$ is the smallest $q$ value measured. This helps to emphasize the mid to high $q$ range.

### 6. Simulation details

In all simulations, the temperature was fixed at 300 K using the v-rescale thermostat with a time constant of 0.5 ps. Coulomb interactions are cut off in real space at 1.0 nm, and we use Particle Mesh Ewald summation with 4th order (cubic) interpolation and a Fourier spacing of 0.12 nm for the long-range electrostatic interactions. The Lennard-Jones interactions were truncated at 1.0 nm and corrected for its effect on energy and pressure [7]. All bonds were constrained using the Lincs algorithm. The integration time step was set to 2 fs and the coordinates of the system were saved every 1000 steps.

The lattice constant of the $H_{II}$-phase was taken from experimental data, determining the simulation box sizes $L_x$ $L_y$ in $x$ and $y$-direction. In $z$-direction, the simulation box size is determined by the ambient pressure. In this $(N, L_x, L_y, P, T)$-ensemble, the pressure is set by coupling the $z$-direction to 1.013 bar using the Berendsen barostat with a compressibility of $4.5\,10^{-5}\,\mathrm{bar}^{-1}$.

The primary challenge for the simulations is to determine the correct number of water molecules in the system. Because the system self-assembles in experiments in excess of water, the chemical potential of water must be identical in the bulk water phase and in the simulated system. The chemical potential is equal to

$$\mu_{water} = k_B T \ln\left(\frac{N_{water}+1}{\langle V\rangle}\Lambda^3\right) + \mu_{ex}, \tag{S37}$$

with $k_B T$ being the thermal energy and $N_{water}/\langle V\rangle$ being the local number of water molecules per unit volume $V$. Because the fugacity $\Lambda$ provides only an arbitrary offset, we set it equal to 1. We calculated the excess chemical potential $\mu_{ex}$ by thermodynamic integration of a water molecule constrained in the center the simulation system by a harmonic potential in $x$ and $y$ directions with a spring constant of $k = 1000\,\mathrm{kJ\,mol^{-1}\,nm^{-2}}$. The integration was performed in two steps over the reaction coordinates $\lambda_1$ and $\lambda_2$, where $\lambda_1 = 0$ is used to interpolate the Lennard-Jones part of the Hamiltonian, and $\lambda_2$ is used to interpolate the Coulomb part [8]. The integrations were performed in 12 steps each using Gaussian integration. The simulation time at each step equaled

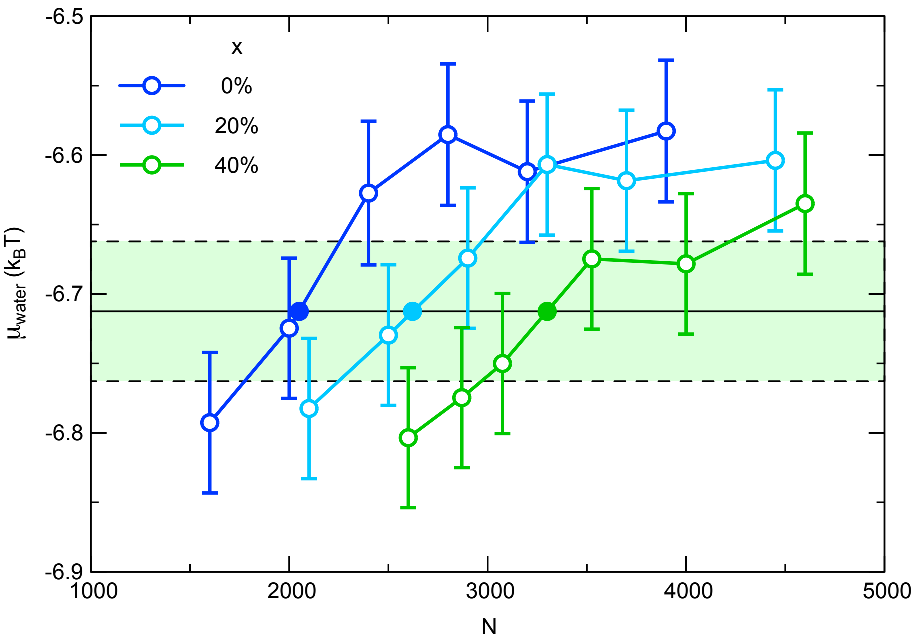


Figure S4: The water chemical potential according to Eq. S37 for pure DOPE (blue symbols) and DOPE: DPhPC mixtures (light blue and green symbols). $x$ denotes the fraction of guest lipid. The black line depicts the chemical potential in bulk water, and the green area depicts its standard deviation. The solid symbols indicate the final values of $N$ used in the simulations.

10 ns and after equilibrating in the $(N, L_x, L_y, P, T)$ ensemble, the simulations were performed in the $(N, V, T)$ ensemble. The number density $N_{water}/\langle V \rangle$ was calculated in a cylinder with a radius of 0.5 nm around the constrained water molecule. The resulting chemical potential of water is shown in Fig. S4 for pure DOPE and its mixture with 40 mol% DPhPC and compared to the bulk value. The final results for the number of water molecules, determined by linear interpolation, equal $N = 2050$ for DOPE in the $\mathrm{H_{II}}$ phase ($x = 0$), $N = 2620$ for DOPE containing 20 mol% DPhPC ($x = 0.2$), and $N = 3300$ for DOPE containing 40 mol% DPhPC ($x = 0.4$).

### 7. Supplementary figures

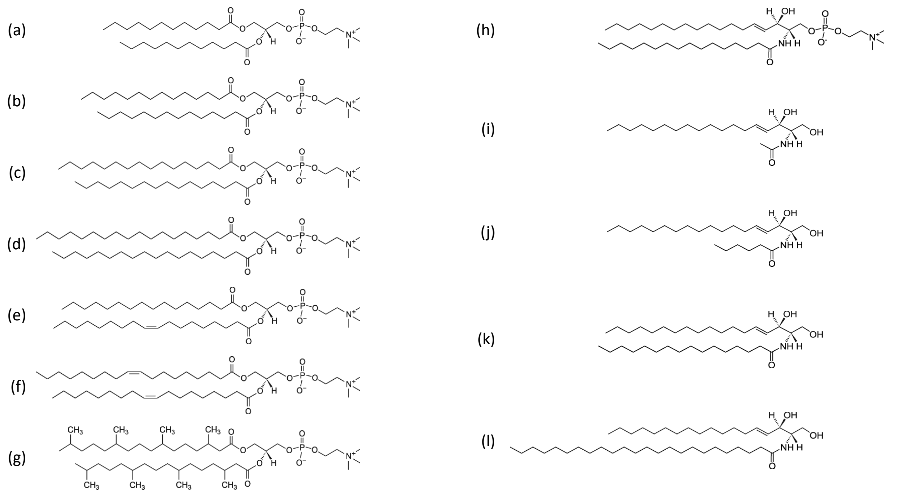


Figure S5: Chemical structure of the studied glycerophospholipids and sphingolipids. (a) 1,2-dilauroyl-*sn*-glycero-3-phosphocholine (DLPC), (b) 1,2-dimyristoyl-*sn*-glycero-3-phosphocholine (DMPC), (c) 1,2-dipalmitoyl-*sn*-glycero-3-phosphocholine (DPPC), (d) 1,2-distearoyl-*sn*-glycero-3-phosphocholine (DSPC), (e) 1-palmitoyl-2-oleoyl-glycero-3-phosphocholine (POPC), (f) 1,2-dioleoyl-*sn*-glycero-3-phosphocholine (DOPC) (g) 1,2-diphytanoyl-*sn*-glycero-3-phosphocholine (DPhPC), (h) N-palmitoyl-D-erythro-sphingosylphosphorylcholine (PSM), (i) N-acetoyl-D-erythro-sphingosine (C2:0 Cer), (j) N-hexanoyl-D-erythro-sphingosine (C6:0 Cer), (k) N-palmitoyl-D-erythro-sphingosine (d18:2(4E,8Z) base) (C16:0 Cer), (l) N-lignoceroyl-D-erythro-sphingosine (d18:2(4E,8Z) base) (C24:0 Cer).

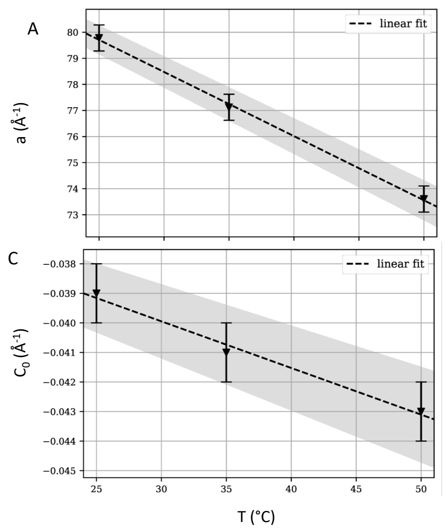


Figure S6: Temperature dependence of the lattice constant (A), lipid area at the neutral plane (B), intrinsic lipid curvature (C) and lipid volume (D) of DOPE.

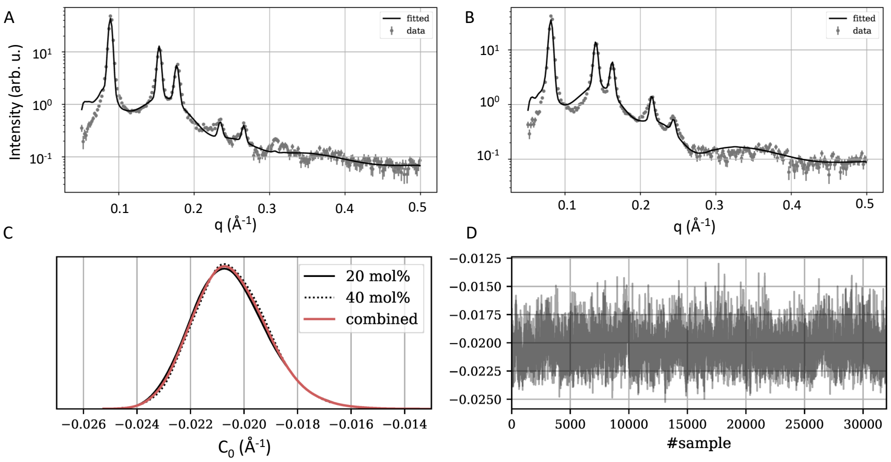


Figure S7: Joint global analysis of DPhPC (20 mol%; panel A and 40 mol%; panel B) in DOPE (35°C). Individual and combined density distribution functions of $C_0$ for both DPhPC concentrations (panel C). Panel D shows the individual MCMC traces.

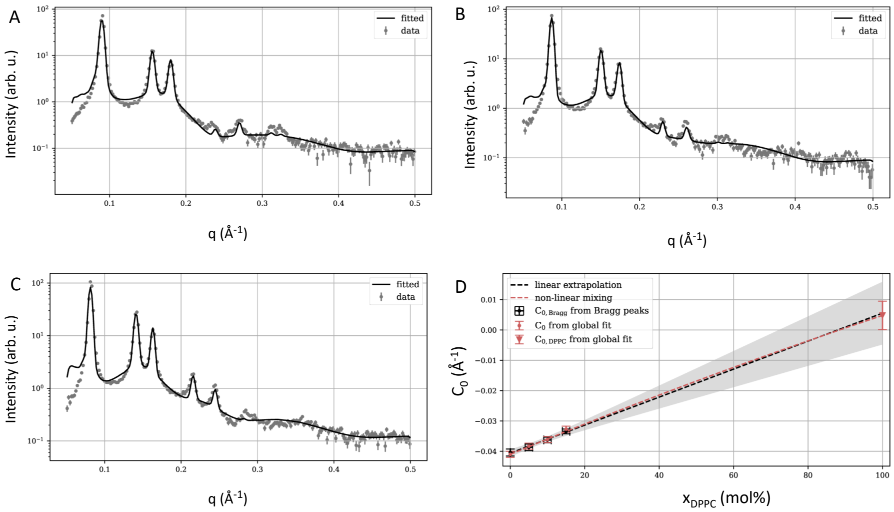


Figure S8: Joint global analysis of DPPC (5 mol%; panel A, 10 mol%; panel B, and 15 mol%) in DOPE (35°C). Panel D compares the total $C_0$ of the system as a function of DPPC concentration as obtained from the global analysis and fitting of the Bragg peaks only [9].

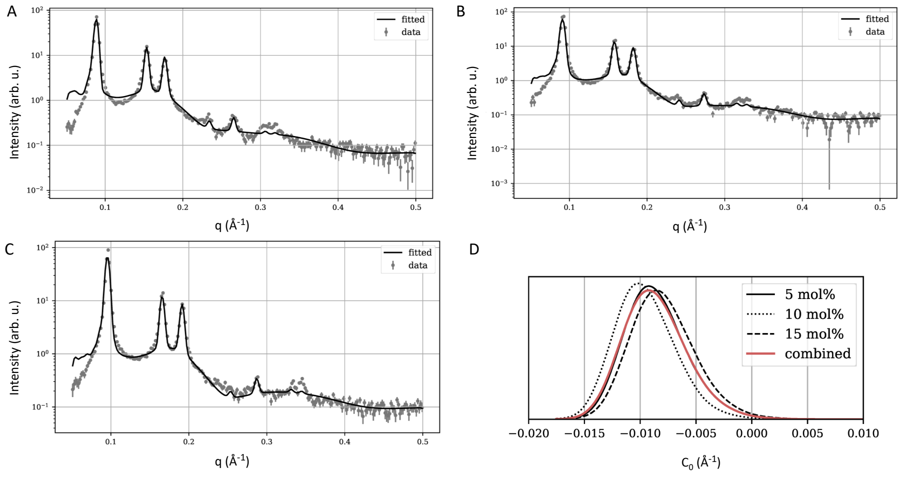


Figure S9: Global analysis of DOPC (5 mol%) at 25°C (panel A), 35°C (panel B) and 50°C (panel C). Panel D shows individual and combined density distribution functions of $C_0$ at 50°C and three different DPPC concentrations.

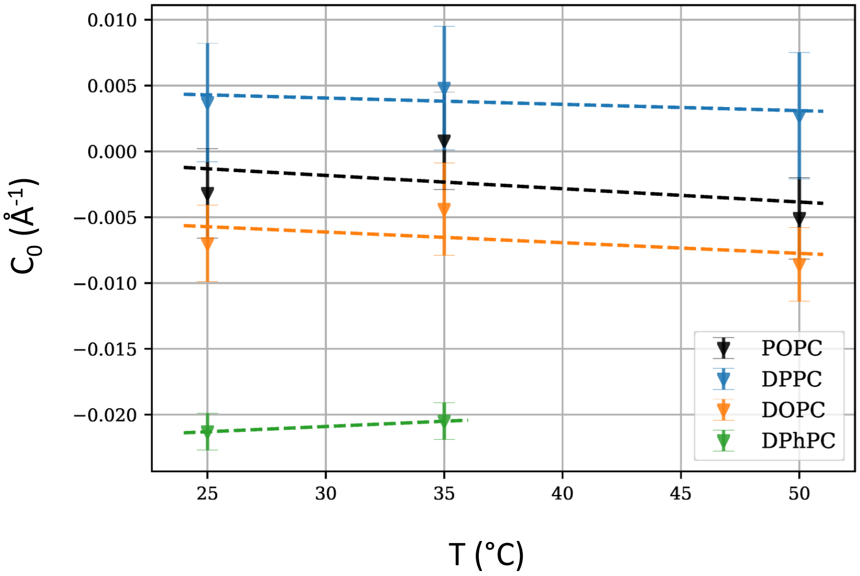


Figure S10: Temperature behavior of $C_0$ of POPC, DPPC, DOPC and DPhPC.

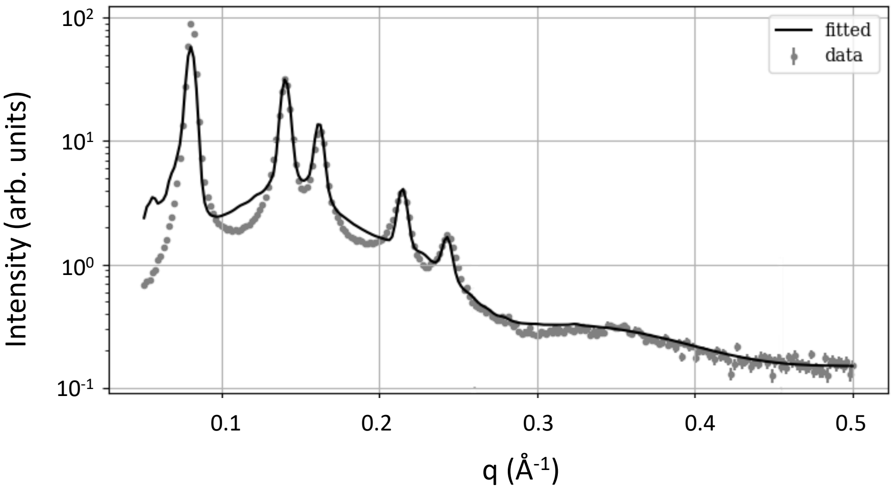


Figure S11: Global analysis of 16:1 PE containing 10 mol% DMPC at 35°C.

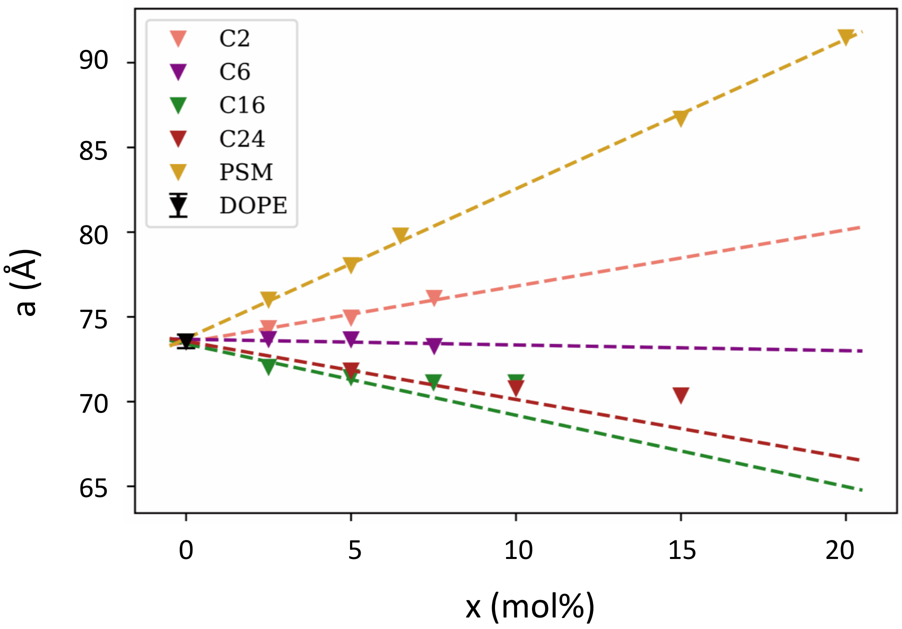


Figure S12: Change of $H_{II}$-lattice constant with sphingolipid concentration at 50°C.

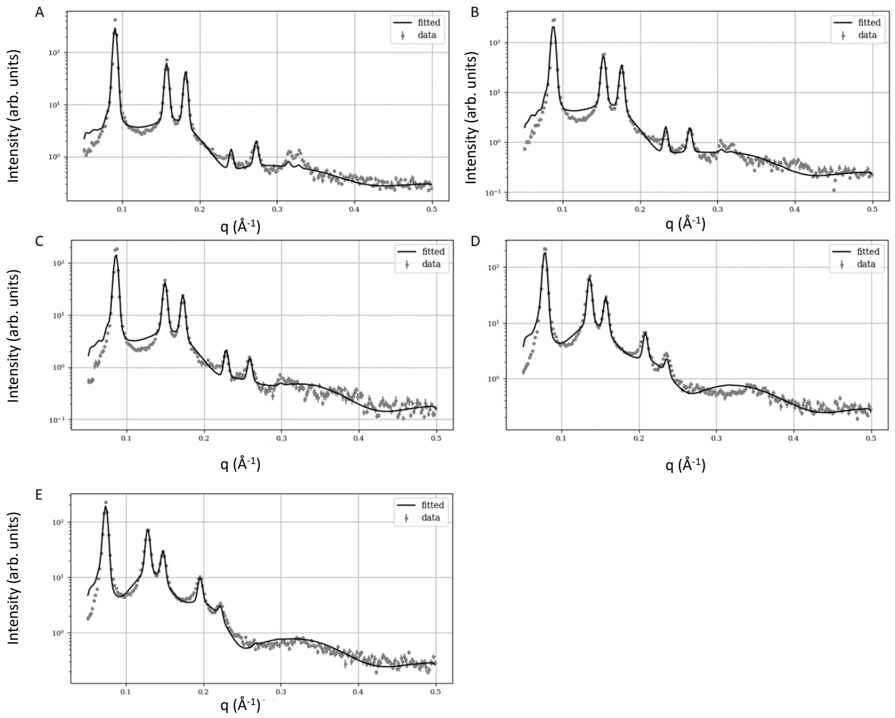


Figure S13: Global analysis of DOPE/PSM mixtures at 35°C. The different PSM concentrations are 2.5% (A), 5.0% (B), 6.5% (C), 15% (D) and 20% (E).

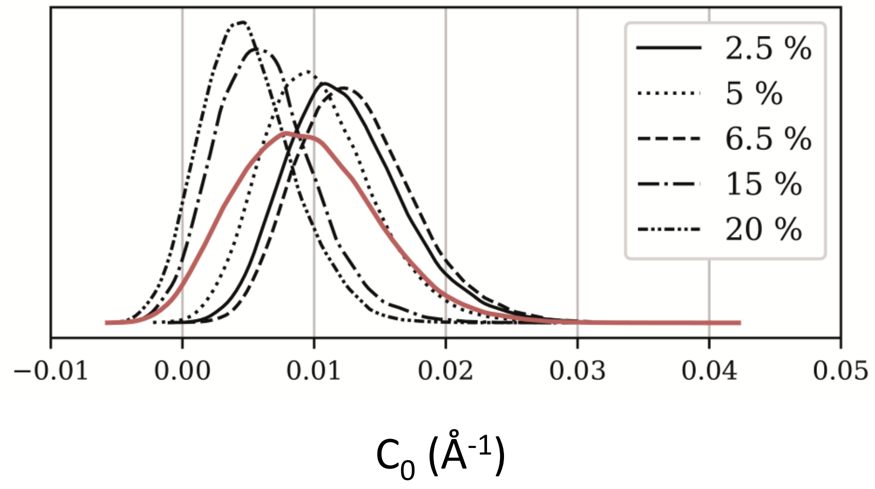


Figure S14: Intrinsic lipid curvature distributions for PSM at different concentrations in DOPE $H_{II}$ templates at 50°C. Distribution centers vary randomly with PSM concentration. The red line gives the average distribution over all PSM concentrations.

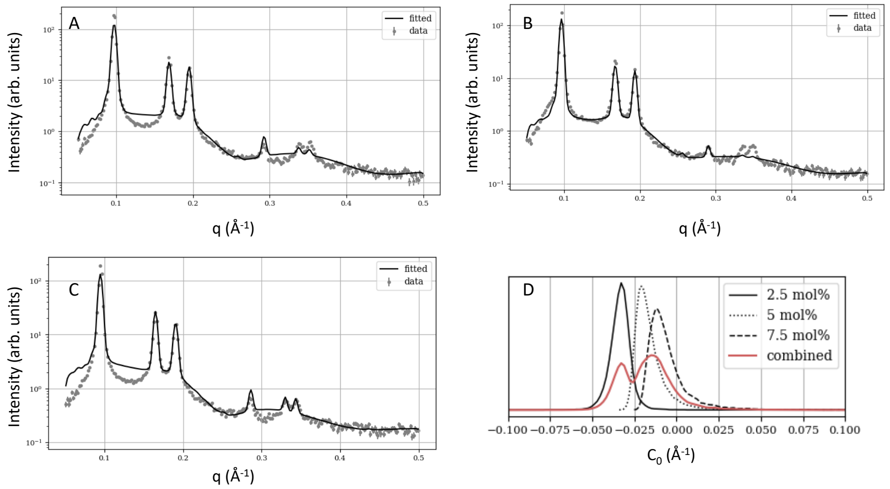


Figure S15: Global analysis of DOPE/Cer2:0 mixtures at 50°C. The different Cer concentrations are 2.5% (A), 5.0% (B), and 7.5% (C). Panel D shows the corresponding intrinsic lipid curvature distributions, where the red solid line represents the average.

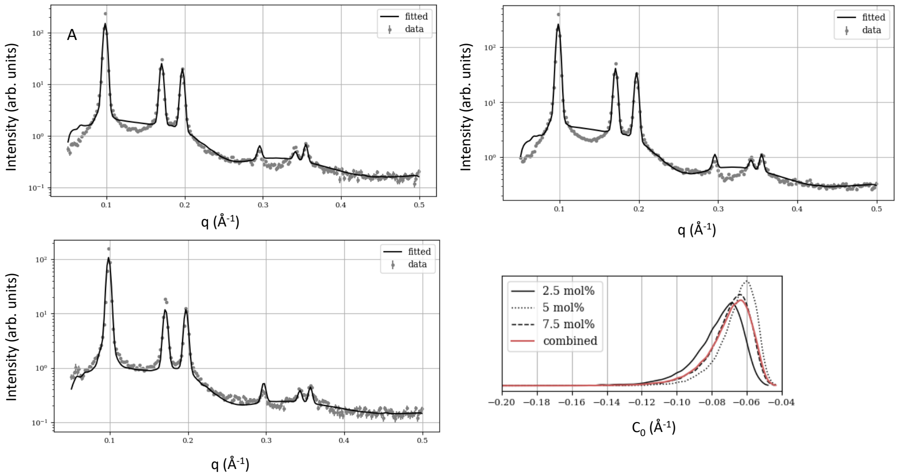


Figure S16: Global analysis of DOPE/Cer6:0 mixtures at 50°C. The different Cer concentrations are 2.5% (A), 5.0% (B), and 7.5% (C). Panel D shows the corresponding intrinsic lipid curvature distributions, where the red solid line represents the average.

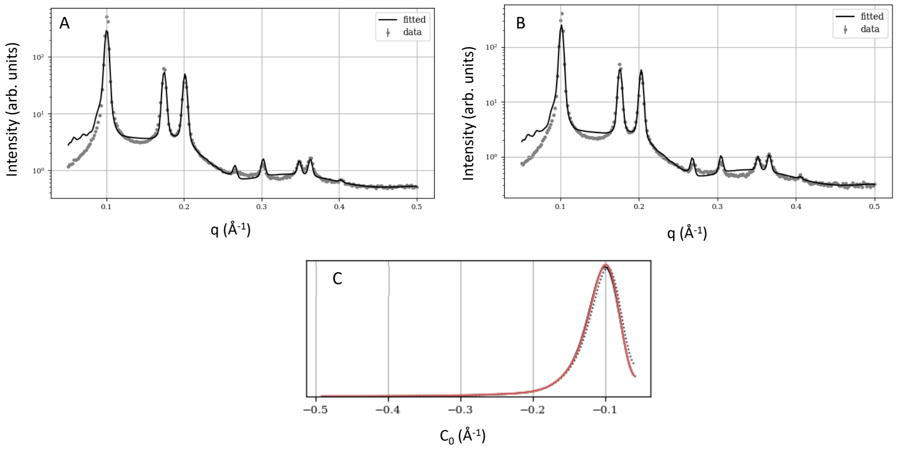


Figure S17: Global analysis of DOPE/Cer16:0 mixtures at 50°C. The different Cer concentrations are 2.5% (A), and 5.0% (B). Panel C shows the corresponding intrinsic lipid curvature distributions, where the red solid line represents the average.

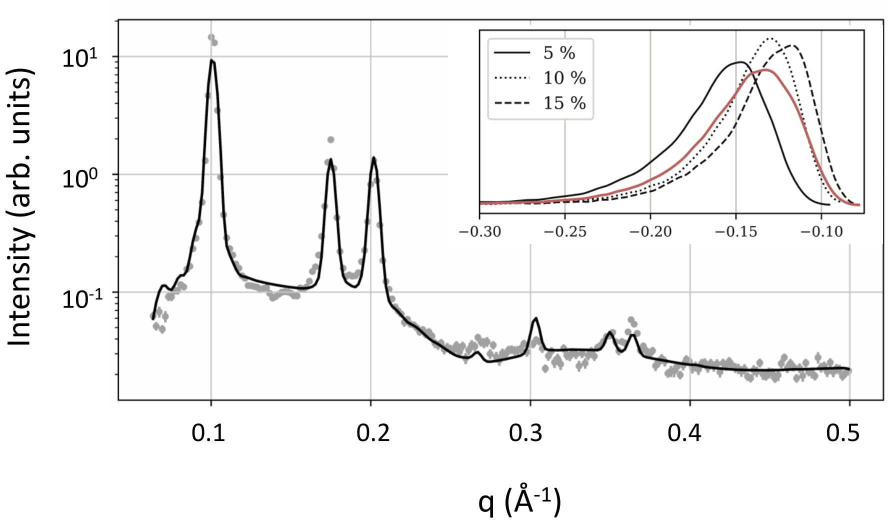


Figure S18: Global analysis of DOPE/Cer24:0 (5 mol%) at 50°C. The insert shows the resulting distributions for $C_0$ including also 10 and 15 mol% Cer24:0.

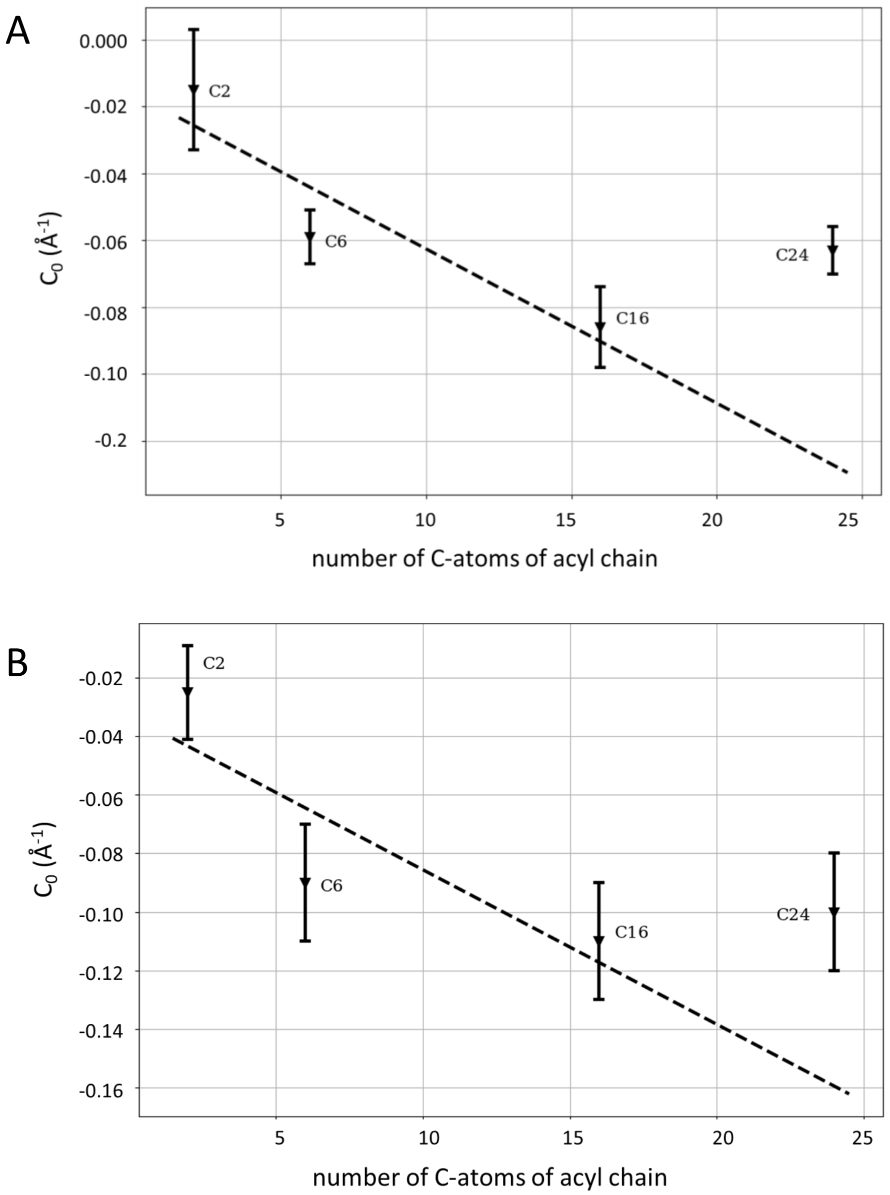


Figure S19: Variation of $C_0$ as a function of chain lenght at 25°C (A) and 35°C.

### 8. Supplementary Tables

Table S1: Overview of adjustable parameters for $H_{II}$ phases composed of single lipids.

| Occurrence | Symbol | Meaning |
|---|---|---|
| Structure factor | $a$ [a] | lattice constant |
| | $\Delta$ | mean square displacement of the lattice points |
| | $n$ | number of hexagonal cells |
| $H_{II}$ form factor | $\sigma_{\text{fluc}}$ | fluctuation constant of lipid unit cell |
| | $C_0$ | intrinsic curvature |
| | $R_W$ | radius of the water core |
| | $V_L$ | lipid volume |
| Monolayer form factor | $c_{\text{mono}}$ | scaling constant |
| | $A_L^{lam}$ | area per lipid |
| Signal scaling | $\Gamma$ | instrumental scaling constant |
| | $I_{\text{inc}}$ | incoherent background |
| | $\gamma$ | scaling constant for intensity error |

[a] derived from Bragg peak positions.

Table S2: Overview of adjustable parameters for lipid mixtures.

| Occurrence | Symbol | Meaning |
|---|---|---|
| Structure factor | $a^a$ | lattice constant |
| | $\Delta$ | mean square displacement of the lattice points |
| | $n$ | number of hexagonal cells |
| $H_{II}$ form factor | $\sigma_{\text{fluc}}$ | fluctuation constant of lipid unit cell |
| | $\xi_{HC}$ | in-plane lipid size ratio at the hydrocarbon interface |
| | $R_W$ | radius of the water core |
| | $V_L$ | lipid volume |
| lamellar form factors | $c_{\text{mono}}$ | monolayer scaling constant |
| | $c_{\text{bi}}$ | bilayer scaling constant |
| | $A_L^{lam}$ | area per lipid |
| | $V_L^{lam}$ | lipid volume in lamellar phases |
| Signal scaling | $\Gamma$ | instrumental scaling constant |
| | $I_{\text{inc}}$ | incoherent background |
| | $\gamma$ | scaling constant for intensity error |

[a] derived from Bragg peak positions.

Table S3: List of externally supplied parameters.

| Parameter | PEs | PCs | PSM | Cer |
|---|---|---|---|---|
| $V_H^L$ (Å$^3$) | $110.25^a$ | $191.98^c$ | $172.62^d$ | $29.56^{df}$ |
| $V_{BB}$ (Å$^3$) | $134.75^a$ | $139.02^c$ | $101.38^d$ | $101.38^d$ |
| $d_{BB}$ (Å) | $4.6^b$ | $4.6^b$ | $4.88^d$ | $4.88^d$ |

[a] from [10]

[b] from [11]

[c] from [12]

[d] from [3]

[e] from [13]

[f] from [4]

Table S4: List of sphingolipid volumina (T= 25/ 35/ 50°C).

| Lipid | $V_L$ (Å$^3$) | $V_L^{lam}$ |
|---|---|---|
| PSM[a] | fitted | 1084.4/1096.9/1156.7 |
| C2:0 Cer[b] | 649.2/647/661.2 | 565.6/577.2/589.9 |
| C6:0 Cer[b] | 757.2/755/769.2 | 673.6/685.2/706.9 |
| C16:0 Cer[b] | 1027.2/1025/1039.2 | 943.6/955.2/976.9 |
| C24:0 Cer[b] | 1243.2/1241/1255.2 | 1159.65/1171.2/1192.9 |

[a] from [3]

[b] derived from [3, 4] and $V_L^{\mathrm{PSM}}$.

Table S5: Temperature dependence of tricosene density, $\tilde{\rho}_{tr}$, molecular volume, $V_{tr}$ and electron density $\rho_{tr}$.

| T (°C) | $\tilde{\rho}_{tr}$ (g/cm$^3$) | $V_{tr}$ (Å$^3$) | $\rho_{tr}$ (Å$^{-3}$) |
|---|---|---|---|
| 15 | 0.8065 | 664.2 | 0.277 |
| 20 | 0.8032 | 667.0 | 0.276 |
| 25 | 0.7999 | 669.7 | 0.275 |
| 30 | 0.7967 | 672.5 | 0.274 |
| 35 | 0.7934 | 675.2 | 0.272 |
| 40 | 0.7901 | 678.0 | 0.271 |
| 45 | 0.7868 | 680.9 | 0.270 |
| 50 | 0.7835 | 683.7 | 0.269 |
| 55 | 0.7803 | 686.6 | 0.268 |
| 60 | 0.7770 | 689.5 | 0.267 |
| 65 | 0.7737 | 692.4 | 0.266 |
| 70 | 0.7705 | 695.3 | 0.265 |
| 75 | 0.7672 | 698.3 | 0.264 |
| 80 | 0.7639 | 701.3 | 0.262 |

Table S6: Temperature dependence of DOPE derived from the global $H_{II}$ data analysis. Bracketed numbers denote the uncertainty of the last digit.

| parameter | 25°C | 35°C | 50°C |
|---|---|---|---|
| $a$ (Å) | 79.8(5) / 80.4[a] | 77.1(5) / 76.9[a] | 73.6(5) / 73.5[a] |
| $\Delta$ (Å$^2$) | 11(2) / 2.9[a] | 9(2) / 3.3[a] | 9(2) /6.7[a] |
| $n$ | 17(2) / 18[a] | 18(3) / 23[a] | 27.8(16) / 29[a] |
| $C_0$ (Å$^{-1}$) | -0.039(1) / -0.039[a] | -0.041(1)/ -0.041[a] | -0.043(1)/ -0.043[a] |
| $V_L$ (Å$^3$) | 1152(11) / 1141[a] | 1154(17) / 1142[a] | 1166(16)/ 1152[a] |
| $R_w$ (Å) | 19.9(6) | 18.3(9) | 17.8(7) |
| $\sigma_{fluc}$ (Å) | 0.159(8) / 0.163[a] | 0.153(18) / 0.163[a] | 0.166(14) / 0.176[a] |
| $c_{mono}$ | 3.4(4) | 2.9(4) | 3.9(5) |
| $A_L^{lam}$ (Å$^2$) | 77(6) / 61.1[a] | 63(5) / 64.9[a] | 88(6) / 65.4[a] |
| $\Gamma$ | 3.8(4) | 3.4(6) | 8.8(12) |
| $I_{inc}$ (arb. u.) | 0.156(4) | 0.148(5) | |
| $d_H$[b] (Å) | 3.2(8) / 4.8[a] | 3.7(11) / 5.2[a] | 3.0(7)/ 5.1[a] |
| $A_0$[b] (Å$^2$) | 58(3) | 59(3) / 62[a] | 63(3) |

[a] from [1]

[b] derived parameter

Table S7: Temperature dependence of 16:1 PE derived from the global $H_{II}$ data analysis. Bracketed numbers denote the uncertainty of the last digit.

| parameter | 25°C | 35°C | 50°C |
|---|---|---|---|
| $a$ (Å) | 80.9(2) | 79.05(1) | 75.5(1) |
| $\Delta$ (Å$^2$) | 2.1(17) | 1.3(12) | 4.6(16) |
| $n$ | 7.5(6) | 8.0(6) | 11(1) |
| $C_0$ (Å$^{-1}$) | -0.0366(3) | -0.0377(3) | -0.0407(5) |
| $V_L$ (Å$^3$) | 1047(9) | 1043.3(9) | 1053(13) |
| $R_w$ (Å) | 20.9(4) | 19.7(4) | 19.0(6) |
| $\sigma_{fluc}$ (Å) | 0.146(5) | 0.146(5) | 0.0153(7) |
| $c_{mono}$ | 3.5(3) | 3.5(3) | 2.8(3) |
| $c_{bi}$ | 4.3(4) | 4.2(4) | 3.4(4) |
| $V_L^{lam}$ (Å$^3$) | 1098(17) | 1104(18) | 1099(19) |
| $A_L^{lam}$ (Å$^2$) | 72(3) | 76(3) | 68(3) |
| $\Gamma$ | 9.8(8) | 10.5(7) | 9.2(8) |
| $I_{inc}$ (arb. u.) | 0.289(5) | 0.289(4) | 0.219(6) |
| ${d_H}^b$ (Å) | 4.3(5) | 4.6(4) | 3.2(7) |
| ${A_0}^b$ (Å$^2$) | 60(2) | 61(2) | 59(2) |

Table S8: Structural parameters of $H_{II}$ phase forming DOPE:DPhPC mixtures at 35°C and increasing DPhPC concentrations. Bracketed numbers denote the uncertainty of the last digits.

| $x$ (%) | $a$ (Å) | $R_W$ (Å) | $R_0$ (Å) | $R_{HC}$ (Å) | $d_H$ (Å) | $V_L$ (Å$^3$) | $\xi_{HC}$ |
|---|---|---|---|---|---|---|---|
| $10^a$ | 78.9 (3) | 18.5 (6) | 26.53 (49) | 28.83 (49) | 5.7 (8) | 1350 (11) | 1.69 (19) |
| $20^b$ | 81.9 (3) | 21.2 (4) | 27.93 (23) | 30.23 (23) | 4.4 (5) | 1365 (30) | 1.38 (4) |
| $40^b$ | 89.2 (3) | 25.5 (4) | 32.68 (32) | 34.98 (32) | 4.9 (5) | 1365 (30) | 1.38 (4) |

$^a$ single analysis

$^b$ joint analysis

Table S9: Structural parameters of $H_{II}$ phase forming DOPE:DPPC mixtures at 35°C and increasing DPPC concentrations. All concentrations were jointly analysed. Bracketed numbers denote the uncertainty of the last digits.

| $x$ (%) | $a$ (Å) | $R_W$ (Å) | $R_0$ (Å) | $R_{HC}$ (Å) | $d_H$ (Å) | $V_L$ (Å$^3$) | $\xi_{HC}$ |
|---|---|---|---|---|---|---|---|
| 5 | 80.6 (3) | 19.6 (4) | 26.11 (14) | 28.41 (14) | 4.2 (4) | 1158 (8) | 1.12 (7) |
| 10 | 83.5 (3) | 21.7 (5) | 27.70 (15) | 30.00 (15) | 3.7 (5) | 1158 (8) | 1.12 (7) |
| 15 | 89.1 (3) | 24.9 (4) | 30.58 (28) | 32.88 (28) | 3.4 (4) | 1158 (8) | 1.12 (7) |

Table S10: Structural parameters of $H_{II}$ phase forming DOPE:PSM mixtures at 35°C and increasing PSM concentrations. All concentrations were jointly analysed. Bracketed numbers denote the uncertainty of the last digits.

| $x$ (%) | $a$ (Å) | $R_W$ (Å) | $R_0$ (Å) | $R_{HC}$ (Å) | $d_H$ (Å) | $V_L$ (Å$^3$) | $\xi_{HC}$ |
|---|---|---|---|---|---|---|---|
| 2.4 (3) | 79.8 (2) | 20.0(3) | 26.32 (7) | 28.63 (7) | 4.2 (3) | 1179 (24) | 1.38 (5) |
| 4.7 (5) | 82.3 (2) | 21.5 (3) | 27.6 (2) | 29.9 (2) | 4.1 (3) | 1179 (24) | 1.38 (5) |
| 6.3 (7) | 83,9 (2) | 23.1 (4) | 28.4 (2) | 30.5 (2) | 3.5 (4) | 1179 (24) | 1.38 (5) |
| 14.9 (15) | 92.1 (2) | 27.8 (4) | 32.7 (3) | 35.0 (3) | 3.5 (4) | 1179 (24) | 1.38 (5) |
| 20.8 (19) | 97.9 (1) | 30.5 (4) | 35.7 (3) | 38.0 (3) | 4.0 (4) | 1179 (24) | 1.38 (5) |

Table S11: Structural parameters of $H_{II}$ phase forming DOPE:C2 mixtures at 50°C and increasing Cer C2 concentrations. All concentrations were jointly analysed. Bracketed numbers denote the uncertainty of the last digits.

| $x$ (%) | $a$ (Å) | $R_W$ (Å) | $R_0$ (Å) | $R_{HC}$ (Å) | $d_H$ (Å) | $V_L$ (Å$^3$) | $\xi_{HC}$ |
|---|---|---|---|---|---|---|---|
| 2.42 (3) | 74.34 (3) | 18.1 (5) | 23.42 (6) | 25.72 (6) | 3.0 (4) | 661 | 0.3 (1) |
| 4.9 (5) | 74.956 (3) | 18.5 (4) | 23.59 (6) | 25.89 (6) | 2.8 (4) | 661 | 0.3 (1) |
| 8.1 (8) | 76.12 (6) | 16.8 (4) | 23.87 (6) | 26.17 (6) | 4.8 (4) | 661 | 0.3 (1) |

Table S12: Structural parameters of $H_{II}$ phase forming DOPE:C6 mixtures at 50°C and increasing Cer C6 concentrations. All concentrations were jointly analysed. Bracketed numbers denote the uncertainty of the last digits.

| $x$ (%) | $a$ (Å) | $R_W$ (Å) | $R_0$ (Å) | $R_{HC}$ (Å) | $d_H$ (Å) | $V_L$ (Å$^3$) | $\xi_{HC}$ |
|---|---|---|---|---|---|---|---|
| 2.5 (3) | 73.695 (3) | 17.8 (4) | 23.09 (6) | 25.39 (6) | 3.1 (4) | 769 | 0.64 (13) |
| 4.9 (5) | 73.68 (2) | 17.6 (2) | 23.0 (1) | 25.3 (1) | 3.2 (3) | 769 | 0.64 (13) |
| 7.1 (8) | 73.28 (4) | 17.8 (5) | 22.8 (1) | 25.1 (1) | 2.8 (4) | 769 | 0.64 (13) |

Table S13: Structural parameters of $H_{II}$ phase forming DOPE:C16 mixtures at 50°C and increasing Cer C16 concentrations. All concentrations were jointly analysed. Bracketed numbers denote the uncertainty of the last digits.

| $x$ (%) | $a$ (Å) | $R_W$ (Å) | $R_0$ (Å) | $R_{HC}$ (Å) | $d_H$ (Å) | $V_L$ (Å$^3$) | $\xi_{HC}$ |
|---|---|---|---|---|---|---|---|
| 2.5 (3) | 72.05 (2) | 16.5 (3) | 21.54 (5) | 23.84 (5) | 2.7 (3) | 1039 | 0.71 (15) |
| 4.8 (5) | 71.45 (2) | 16.2 (2) | 21.15 (9) | 23.45 (9) | 2.9 (3) | 1039 | 0.71 (15) |

Table S14: Structural parameters of $H_{II}$ phase forming DOPE:C24 mixtures at 50°C and increasing Cer C24 concentrations. All concentrations were jointly analysed. Bracketed numbers denote the uncertainty of the last digits.

| $x$ (%) | $a$ (Å) | $R_W$ (Å) | $R_0$ (Å) | $R_{HC}$ (Å) | $d_H$ (Å) | $V_L$ (Å$^3$) | $\xi_{HC}$ |
|---|---|---|---|---|---|---|---|
| 5.0 (5) | 71.844 (2) | 16.9 (4) | 21.6 (2) | 23.6 (2) | 2.8 (4) | 1255 | 0.6 (2) |
